# Selective bHLH relay factors and modular enhancers decode Notch signaling during neuronal diversification in the *Drosophila* medulla

**DOI:** 10.64898/2026.06.05.730496

**Authors:** Yu Zhang, Tejus Sreelal, Hailun Zhu, Raymond Warner Jones, Michelle Seby, Xin Li

**Affiliations:** Department of Cell and Developmental Biology, University of Illinois Urbana-Champaign, Urbana, IL 61801-3761

**Author notes:** Equal contribution (Yu Zhang, Tejus Sreelal).

## Abstract

A central question in developmental biology is how a single signaling pathway can generate diverse developmental outcomes. One prominent example is Notch-dependent binary fate choice, which is reiteratively utilized during development to sequentially generate multiple distinct pairs of cell fates. To investigate the transcriptional mechanisms that diversify Notch signaling outputs, we profiled gene expression and chromatin accessibility simultaneously in single cells of the developing *Drosophila* medulla, identified and analyzed the cis-regulatory enhancer elements (CREs) for representative neuronal transcription factor (nTF) genes, whose expression depends on the Notch status. Canonical models predict that Notch target genes of the Hey/Hes family act primarily as transcriptional repressors and bind the CACGTG E-box motif to suppress Notch-off transcriptional programs in Notch-on cells. Contrary to the prediction, we found that Hey is not required for repression of Notch-off nTF genes. Instead, Hey is required for activation of a subset of Notch-on nTF genes. Using the Notch-on nTF gene *bsh* as an example, we identified a CRE that recapitulates the endogenous expression pattern and demonstrated that a single-base-pair mutation in a CACGTG type E-box present in the CRE abolishes its enhancer activity. We further identified additional relay factors that mediate Notch-dependent transcriptional outputs. The bHLH factors Sim/Tgo activate a Notch-on nTF through binding to the AACGTG variant E-boxes within its CRE, whereas the bHLH factor Tap activates a Notch-off nTF through a CRE containing different types of E-boxes. Together, these findings reveal that Notch signaling is not relayed through a single universal downstream effector. Instead, distinct bHLH factors decode Notch status through different classes of E-box motifs embedded within target enhancers. Finally, we show that enhancer architecture is modular, allowing temporal identity and Notch-status information to be integrated through the same or distinct CREs to generate precise patterns of nTF expression. We propose that diversification of Notch-dependent cell fates arises through a modular transcriptional relay and enhancer-decoding mechanism in which multiple bHLH factors act on distinct E-box motifs to convert a common signaling input into diverse developmental outcomes.

## INTRODUCTION

Notch (N) signaling pathway is a highly conserved signaling pathway with pleiotropic and essential roles in controlling developmental decisions^1,2^. One major mechanism by which Notch signaling controls cell fates is through Notch-dependent binary fate choice, a strategy that is often used reiteratively during development to generate sequential cell fate decisions. Particularly in the nervous system, Notch-dependent binary fate choice can substantially expand the neural diversity produced by the temporal and spatial patterning of neural progenitors^3–9^. Notch-dependent binary fate choice was first discovered in *Drosophila* Sensory Organ Precursor (SOP) and neuroblast (NB) lineages three decades ago^10,11^. *Drosophila* type I NBs undergo repeated asymmetric divisions to self-renew while generating a series of intermediate progenitors known as Ganglion Mother Cells (GMCs). Each GMC subsequently divides once to produce two postmitotic progeny that typically adopt distinct fates. Both asymmetric divisions involve Notch signaling. Here we focus on the GMC division that generates two distinct postmitotic daughter cells. Numb, a negative regulator of Notch signaling, is asymmetrically segregated into one of the two daughters. The daughter cell that inherits Numb will become Notch-off (N-off), while the sibling cell will have active Notch signaling and adopt the N-on fate ^10,11^. Subsequent studies demonstrated that Notch-dependent binary fate choice is extensively used throughout invertebrate nervous systems to diversify neuronal identities^3–9^. Within a lineage, all N-on progeny collectively form the Notch-on hemilineage, whereas all N-off progeny constitute the N-off hemilineage. These two hemilineages often exhibit striking differences in neuronal morphology, connectivity, and neurotransmitter expression^3–9^. More recent studies in vertebrate systems have further demonstrated the conservation of Notch-dependent binary fate choice in the diversification of neural fates ^12–15^.

The core components of the Notch pathway have been extensively characterized^16,17^. Upon ligand binding to the extracellular domain of the Notch receptor, the Notch receptor undergoes sequential proteolytic cleavages by a metalloprotease and γ-secretase, releasing the Notch intracellular domain (NICD). NICD subsequently translocates into the nucleus, where it binds with the ubiquitous transcription factor CBF-1/suppressor of hairless/Lag1 (CSL, also called recombination signal binding protein-J, RBPJ). Together, they recruit the transcriptional coactivator Mastermind to activate the expression of Notch target genes, including members of the Hes (Hairy/Enhancer of split) and Hey (Hairy/E(spl)-related with YRPW motif) gene families^16,17^. These Hey and Hes genes encode highly conserved and closely related basic helix-loop-helix (bHLH) transcription factors. bHLH factors bind DNA as dimers to different variants of E-boxes “CANNTG”, with each factor contacting a half site (CAN) on opposite strands ^18^. The consensus binding motif for Hey and Hes proteins is the Hey type E-box (CACGTG), which is highly conserved in invertebrates and vertebrates^18^. Hey and Hes proteins have been generally considered to function primarily as transcriptional repressors^17,19,20^. Despite extensive knowledge of the Notch signaling machinery and its target genes, the mechanisms by which these targets mediate Notch-dependent fate divergence remain poorly understood. It is also not clear how a single signaling pathway instructs multiple pairs of neural fates within an individual neuroblast lineage.

We address these questions in the *Drosophila* medulla, part of the visual processing center in the brain called the optic lobe. During development, medulla neuroblasts generate more than 100 different types of medulla neurons through a combination of temporal patterning, spatial patterning and Notch-dependent binary fate choice^5,21,22,6,23–25^. Medulla neuroblasts are temporally patterned by the sequential expression of a cascade of Temporal Transcription Factors (TTFs), including Homothorax (Hth), SoxNeuro (SoxN), doublesex-Mab related 99B (Dmrt99B), Odd paired (Opa), Earmuff (Erm), Eyeless (Ey), Homeobrain (Hbn), Sloppy paired 1 and 2 (Slp), Scarecrow (Scro), Dichaete (D), BarH1 and BarH2, Tailless (Tll) and Glial cells missing (Gcm)^5,21,24,25^. In addition, the neuroepithelium that gives rise to medulla neuroblasts is spatially patterned to allow generation of specialized neural types^22,26,27,8^. Notch-dependent binary fate choice further doubles the neural diversity generated by temporal and spatial patterning. Immature Notch-on neurons express Hey^26,28^, the sole ortholog of vertebrate Hey proteins including Hey1, Hey2 and HeyL^29^, followed by the expression of a TF called Apterous (Ap) in all Notch-on neurons ^5,26,28^. The N-on and N-off hemilineage neurons have different morphology, connectivity and neurotransmitter expression^6,28,9,8^. Different neuronal transcription factors (nTFs) are expressed in N-on and N-off hemilineages, and some of them have been confirmed to control neuron fate, morphology and neurotransmitter expression^30,31,6,32,33,24,25^. To investigate how Notch signaling regulates the expression of these nTFs to control the divergent fates, we profiled gene expression and chromatin accessibility simultaneously at single-cell level in the developing medulla, compared enriched motifs in differentially accessible peaks between N-on and N-off neurons, and systematically cloned and analyzed the cis-regulatory enhancer elements (CREs) controlling the expression of representative Notch-on or Notch-off nTFs. Opposite to the generally accepted transcriptional repressor role of Notch target gene Hey, our results demonstrate that Hey is required for activating the expression of certain N-on nTF genes. Using the Notch-on nTF gene *bsh* as an example, we identified a CRE that recapitulates the endogenous expression pattern and demonstrated that a single-base-pair mutation in a CACGTG type E-box abolishes enhancer activity. Furthermore, our study identified additional bHLH family relay factors of Notch signaling, including Sim/Tgo, which are required to activate another N-on nTF gene through a variant E-box “AACGTG” present in its CRE; and Tap, which is required to activate the expression of a N-off nTF. Our results also demonstrate the modularity of enhancers, and that temporal identity and Notch-status information can be integrated on the same or different CREs to generate precise patterns of nTF expression. Together, these findings reveal how a common Notch signaling input is transformed into diverse developmental outcomes through the coordinated action of multiple bHLH relay factors and modular enhancer architectures.

## RESULTS

### Single-cell multiome and scATAC-seq analyses identify Notch-status-associated nTF programs and candidate regulatory elements

Notch signaling mediates the binary fate choice between sibling neurons generated by medulla GMCs (Figure 1A). Twin-spot MARCM studies have shown that Ap distinguishes the two sibling neurons born from the same GMC, with the Notch-on one expressing Ap and the Notch-off counterpart lacking Ap ^5^. Previous studies also showed that Notch signaling regulates selected identity factors in medulla neurons, including Ap, Ey and Toy ^5^. However, whether and how canonical Notch signaling broadly organizes nTF programs across temporal contexts, and how these diverse outputs are encoded at the cis-regulatory level, remain unclear.

**Figure 1.**
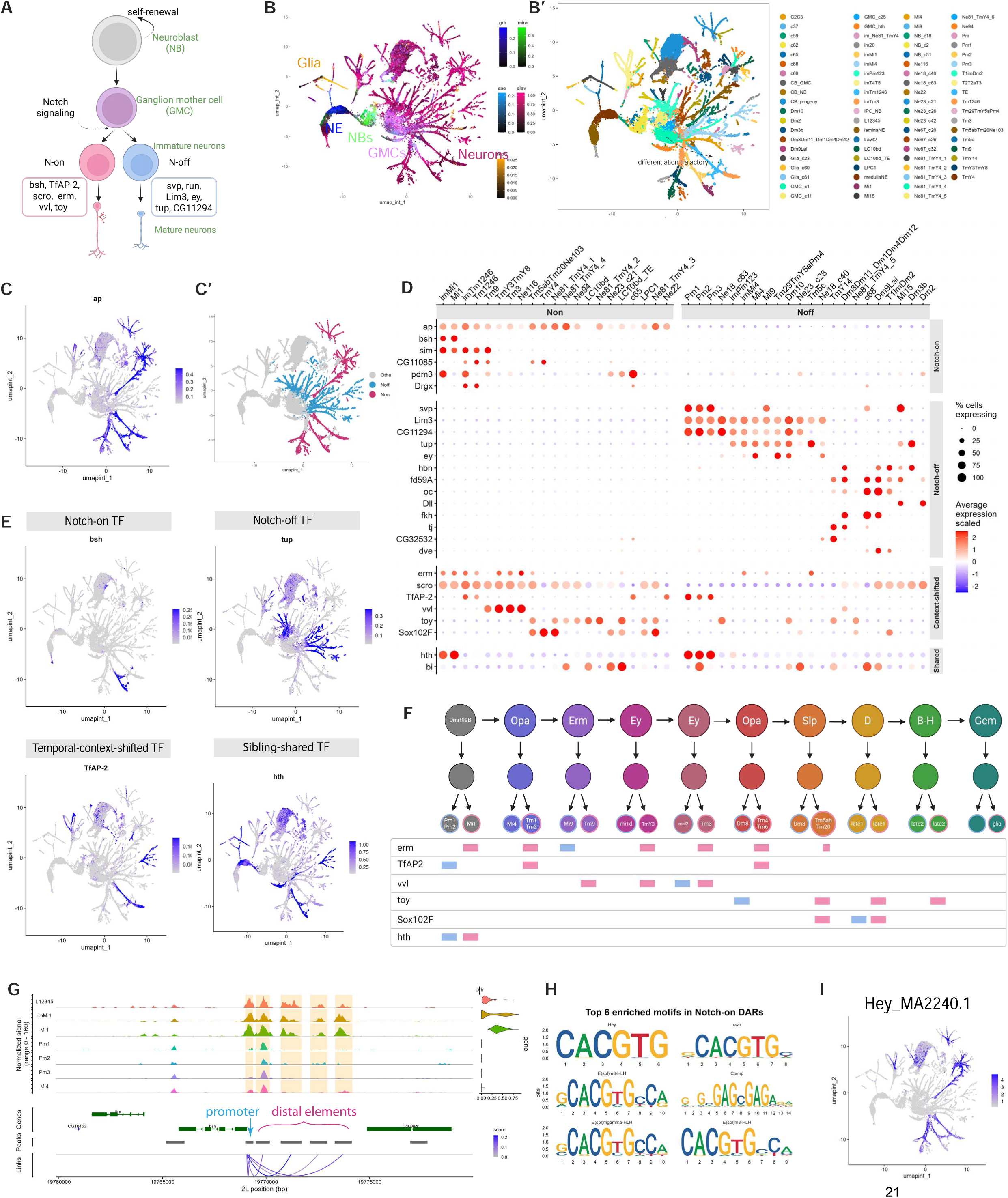
Single-cell multiomics identifies Notch-status-associated nTF programs and candidate cis-regulatory landscapes. (A) Schematic of Notch-dependent binary fate choice during medulla neuron development. Each medulla ganglion mother cell (GMC) generates a pair of sibling neurons with different Notch status: a Notch-on sibling and a Notch-off sibling. (B) UMAP of the scATAC-seq dataset annotated with major developmental cell states, including neuroepithelial cells, neuroblasts, GMCs, glia, and neurons. (B′) Detailed annotation of the scATAC-seq UMAP. The dashed arrow indicates the inferred differentiation trajectory from neuroblasts to GMCs and differentiating neurons. (C) Feature plot showing ap expression, which marks Notch-on neuronal populations. (C′) UMAP showing assignment of neuronal populations into Notch-on and Notch-off groups based on ap expression and cluster identity. (D) Dot plot showing expression of transcription factors classified according to their relationship with Notch status and sibling-pair logic. TFs are grouped into Notch-on sibling-restricted, Notch-off sibling-restricted, temporal-context-shifted, and sibling-shared classes. Within each Notch-status group, clusters are arranged according to relative birth order, with early-born populations on the left and late-born populations on the right. (E) Feature plots showing representative TFs from each class: bsh as a Notch-on sibling-restricted TF, tup as a Notch-off sibling-restricted TF, TfAP-2 as a temporal-context-shifted TF, and hth as a sibling-shared TF. (F) Schematic summarizing the relationship between neuroblast temporal origin, sibling-pair identity, and expression of temporal-context-shifted and sibling-shared TFs. TFs such as TfAP-2, erm, vvl, toy, and Sox102F can appear in both Notch-status populations across the dataset because they are expressed in progeny generated at different temporal windows, whereas hth represents a true sibling-shared factor. (G) Genome browser coverage plot showing chromatin accessibility and peak-gene links at a representative nTF locus. Distal accessible elements, rather than broadly accessible promoters, show restricted accessibility patterns and are linked to nTF expression. (H) Top enriched motifs in Notch-on-associated differentially accessible regions compared with Notch-off-associated regions. Enriched motifs include E-box-like motifs recognized by bHLH-family transcription factors. (I) Feature plot showing chromVAR deviation scores for the Hey-class/E-box-like motif Hey_MA2240.1, highlighting enrichment of this motif activity in Notch-on-associated populations.

To define how Notch status is associated with nTF expression, we generated and analyzed paired single-cell multiome data from developing third-instar larval optic lobes using the 10x Genomics platform. After quality control, both the RNA modality and the ATAC modality resolved clean separated clusters (6,622 cells; Figure S1A). We annotated multiome clusters using the RNA modality, taking advantage of well-established transcriptome-based atlas of medulla cell states^23,25^. In parallel, we generated a larger scATAC-seq dataset to resolve medulla cell states and candidate regulatory elements with greater granularity (36,076 cells). We integrated the larger scATAC-seq dataset with the ATAC modality of the annotated multiome dataset and transferred cell-state annotations to the larger accessibility map (Figure 1B-B’). Because RNA-only, ATAC-only, and integrated analyses produced different UMAP layouts, we used the integrated ATAC-based UMAP throughout the paper to ensure consistent visualization across datasets and analyses. The resulting UMAP showed an ordered progression from neuroblasts to GMCs, immature neurons, and more differentiated neuronal populations, consistent with the previously described medulla differentiation trajectory^23,25^. We therefore used this trajectory as a reference to distinguish immature and more mature neuronal states in downstream analyses. Since Ap marks the Notch-on sibling, we used *ap* expression to distinguish Notch-on from corresponding Notch-off neuronal populations (Figure 1C-C’).

We next examined transcription factor expression across Notch-on and Notch-off populations. This analysis revealed four major classes of nTFs based on the relationship of their expression patterns and Notch-status. First, a small set of TFs, including *ap*, *bsh*, *sim*, *CG11085*, *pdm3*, and *drgx*, were expressed exclusively in Notch-on neurons. Second, several TFs, including *svp*, *Lim3*, *CG11294*, *tup*, *ey*, *hbn*, *fd59A*, *oc*, *dll*, *fkh*, *tj*, *CG32532*, and *dve*, were restricted to Notch-off siblings. Third, TFs such as *erm*, *scro*, *TfAP-2*, *vvl*, *toy*, and *Sox102F* appeared to be expressed in both Notch-on and Notch-off populations when neurons generated across all temporal stages were analyzed together (Figure 1D-E). However, this apparent sharing usually reflected expression in progeny generated at different neuroblast temporal windows, rather than expression in both siblings from a single GMC division(Figure 1F). We refer to this class as temporal-context-shifted TFs. Finally, a small number of TFs, such as *hth* and *bi*, were expressed in both siblings and therefore represented true sibling-shared factors (Figures 1D-1F). This classification distinguishes global Notch-status sharing from true sibling-level sharing and shows that nTF expression reflects the intersection of Notch status with neuroblast temporal origin.

We next asked how these selective nTF expression patterns are encoded at the cis-regulatory level. Consistent with previous large-scale cis-regulatory analyses of the adult *Drosophila* brain ^34^, chromatin accessibility analysis of the developing optic lobe revealed that nTF promoters were broadly accessible across medulla cell states (Figure S1B), whereas distal accessible elements showed more restricted patterns (Figure 1G). We therefore focused on distal elements as candidate enhancers. The paired multiome data enabled peak-gene linkage between accessible elements and nTF expression, while the larger scATAC-seq dataset provided higher-resolution accessibility patterns across cell states. Together, these analyses nominated candidate regulatory elements near nTF loci, including *bsh*, *TfAP-2*, *tup*, and *Lim3* (Figures 1G; Supplementary Table 1). These linked elements provided candidate enhancer modules for functional analysis in subsequent sections.

To identify the transcriptional regulators associated with Notch-dependent fate divergence, we compared chromatin accessibility profiles between Notch-on and Notch-off hemilineages. Using the top differentially accessible peaks between the two populations, we performed motif enrichment analysis to identify Notch-status-associated regulatory signatures. The most significantly enriched motif in chromatin regions preferentially accessible in Notch-on neurons was the canonical E-box sequence, CACGTG (Figure 1H; Supplementary Table 2). To determine whether this sequence enrichment was accompanied by increased accessibility at motif-containing elements within Notch-on cells, we next calculated chromVAR deviation scores. ChromVAR estimates, for each cell, whether peaks containing a given motif are more accessible than expected relative to matched background peaks. Consistent with the enrichment analysis, E-box motifs recognized by multiple basic helix-loop-helix (bHLH) factors, including E(spl) family members and Hey, showed elevated deviation scores in Notch-on populations (Figure 1I and Figure S1C). These results suggest that Notch-on neurons exhibit increased accessibility at CACGTG type E-box-containing regulatory elements, implicating bHLH factors binding to this motif as candidate mediators of Notch-dependent regulatory divergence.

Together, these analyses established the framework for the study. Paired multiome data linked nTF expression with candidate regulatory elements, while the larger scATAC-seq dataset provided higher-resolution accessibility and motif information across medulla cell states. These datasets revealed that Notch status is associated with nTF expression in a temporal-context-dependent manner and nominated candidate enhancer modules and E-box-related motif signatures for functional testing.

### Canonical Notch signaling establishes Notch-status-specific nTF expression

Having identified Notch-status-associated nTF programs, we next tested whether these programs depend on canonical Notch signaling (Figure 2A). Previous studies established that Notch signaling mediates the binary fate decision between sibling neurons generated from medulla GMC divisions, but only a limited number of downstream transcription factors, including Ap, had been examined. To define the broader role of Notch signaling in medulla neuronal diversification, we perturbed *Suppressor of Hairless* (*Su(H)*), the central transcriptional effector of canonical Notch signaling, using mosaic RNAi clones.

**Figure 2.**
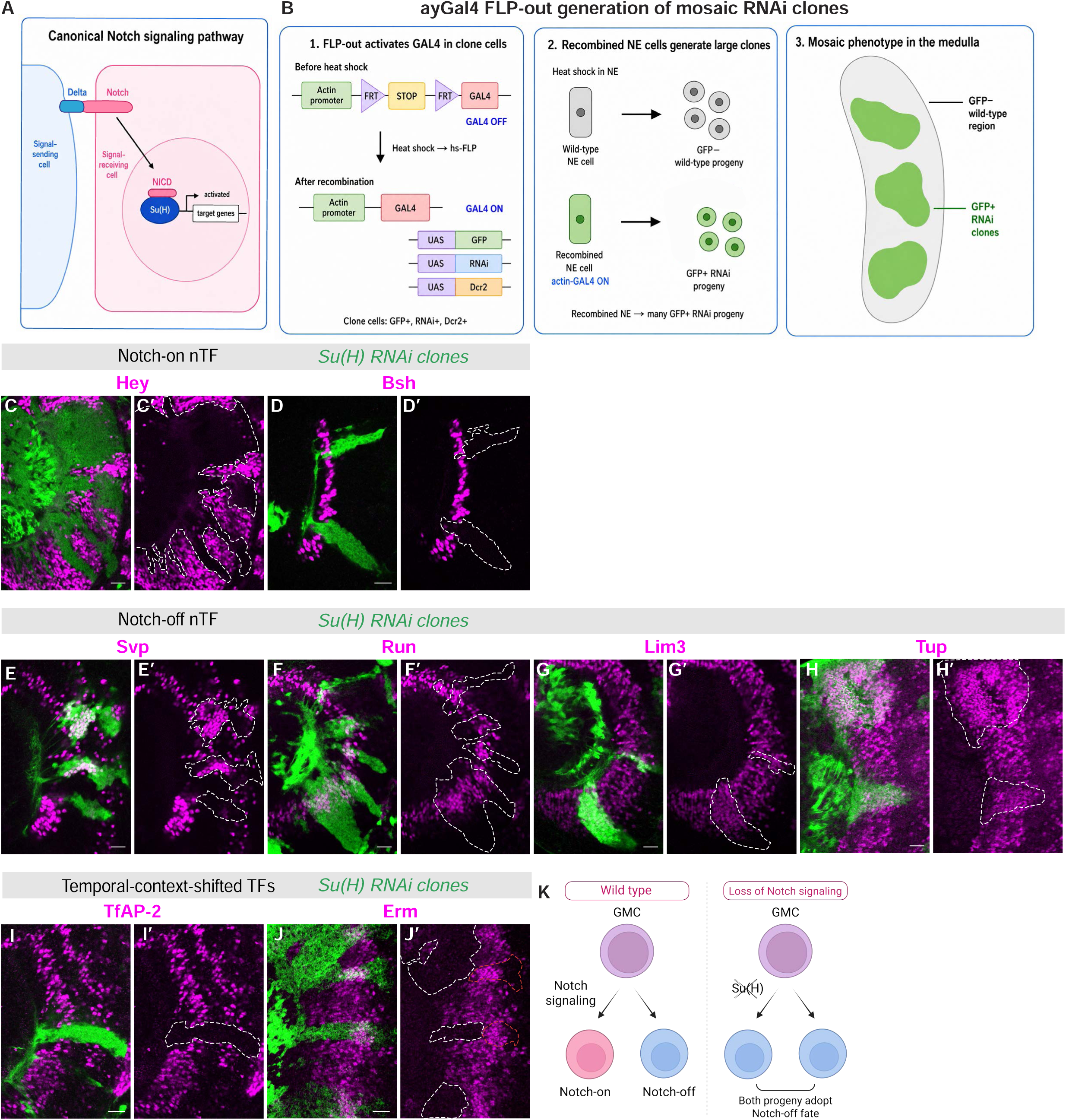
Canonical Notch signaling broadly controls Notch-dependent nTF programs. (A) Schematic of the canonical Notch signaling pathway. Ligand-dependent Notch activation releases NICD, which acts through Su(H) to regulate target gene expression. (B) Strategy for generating mosaic RNAi clones using the ayGal4 FLP-out system in the developing Drosophila medulla. Heat-shock-induced FLP recombination removes the STOP cassette, activating Gal4 and driving expression of UAS-GFP, UAS-RNAi, and UAS-Dcr2 in clone cells. GFP marks RNAi-expressing clones. (C and D) Notch-on nTF expression is lost in *Su(H) RNAi* clones. Hey expression is abolished in *Su(H)* RNAi clones (C–C′), and Bsh expression is lost in clone cells (D–D′). (E–H) Notch-off nTF expression expands ectopically in *Su(H)* RNAi clones. Svp (E–E′), Run (F–F′), Lim3 (G–G′), and Tup (H–H′) expand into clone regions. (I and J) Temporal-context-shifted TFs show domain-specific responses after loss of Su(H). TfAP-2 expression in Notch-on cells is lost in *Su(H)* RNAi clones (I–I′), whereas Erm expression is reduced in the Notch-on domain and expands in the Notch-off domain (J–J′). White dashed outlines indicate Notch-on clone regions, and red dashed outlines indicate Notch-off clone regions. (K) Model summarizing the effect of Su(H) loss. In wild type, Notch signaling promotes Notch-on identity while restricting Notch-off programs. In the absence of canonical Notch signaling, Notch-on nTFs are lost and both sibling progeny adopt Notch-off-like transcriptional features. Dashed outlines mark the RNAi clone regions. Scale bars, 20 μm.

We generated RNAi clones using the *ayGal4* FLP-out system^35^, in which heat-shock-induced FLP excises a STOP cassette to activate *actin*-Gal4-driven RNAi expression. This allowed us to induce *Su(H)* RNAi clones and assess nTF expression cell-autonomously in developing medulla lineages (Figure 2B).

Loss of *Su(H)* abolished Hey expression (Figure 2C,C’), consistent with Hey acting downstream of canonical Notch signaling. *Su(H)* RNAi also broadly disrupted Notch-status-specific nTF expression, with three classes of effects that reflected the Notch-status and temporal organization defined in Figure 1D. First, Notch-on-restricted nTF expression, including Bsh, was lost (Figure 2D,D’). Second, Notch-off-restricted factors, including Svp, Run, Lim3, Tup and Ey, expanded ectopically to the siblings and became compact (Figure 2E-H’). Third, temporal-context-shifted TFs, such as Erm and TfAP-2, showed domain-specific responses: their Notch-on expression domains were reduced or lost, whereas their Notch-off expression domains were preserved or expanded (Figures 2I–J’).

These reciprocal and domain-specific changes indicate that canonical Notch signaling broadly promotes Notch-on nTF expression while preventing inappropriate expansion of Notch-off nTF programs across multiple temporal contexts (Figure 2K). Together, these findings suggest that Su(H)is required for the Notch-status input, which is interpreted in the context of temporal identity. Because several candidate relays downstream of Notch are bHLH factors that recognize related E-box-like motifs, we next asked whether specific E-box-binding relay factors link canonical Notch signaling to selected nTF outputs.

### Hey is required to activate the expression of selected Notch-on nTFs

To evaluate Hey as a candidate Notch-on relay, we first examined its expression pattern. Hey is a canonical Notch-responsive bHLH transcription factor, and our previous work showed that Hey is transiently expressed in immature Notch-on neurons^28^. Unlike many Notch-on nTFs that are restricted to specific neuron types, Hey showed a broad Notch-on pattern similar to Ap, suggesting that it could act as a general relay for Notch-on identity.

In the single-cell RNA data, however, Hey transcripts were detected earlier than ap transcripts and were present in immature neurons from both Notch-status groups (Figures 3A and 1C). To define the protein expression pattern, we stained wild-type brains using anti-Hey together with HCR FISH probe set against *ap mRNA*. Hey protein appeared at the same developmental stage as ap transcripts and was restricted to Ap+ Notch-on neurons (Figures 3B–B’2). Thus, although hey transcripts were detected earlier and more broadly, Hey protein was Notch-on restricted. This discrepancy suggests that Hey Notch-status specificity is refined beyond transcription, potentially through post-transcriptional regulation.

**Figure 3.**
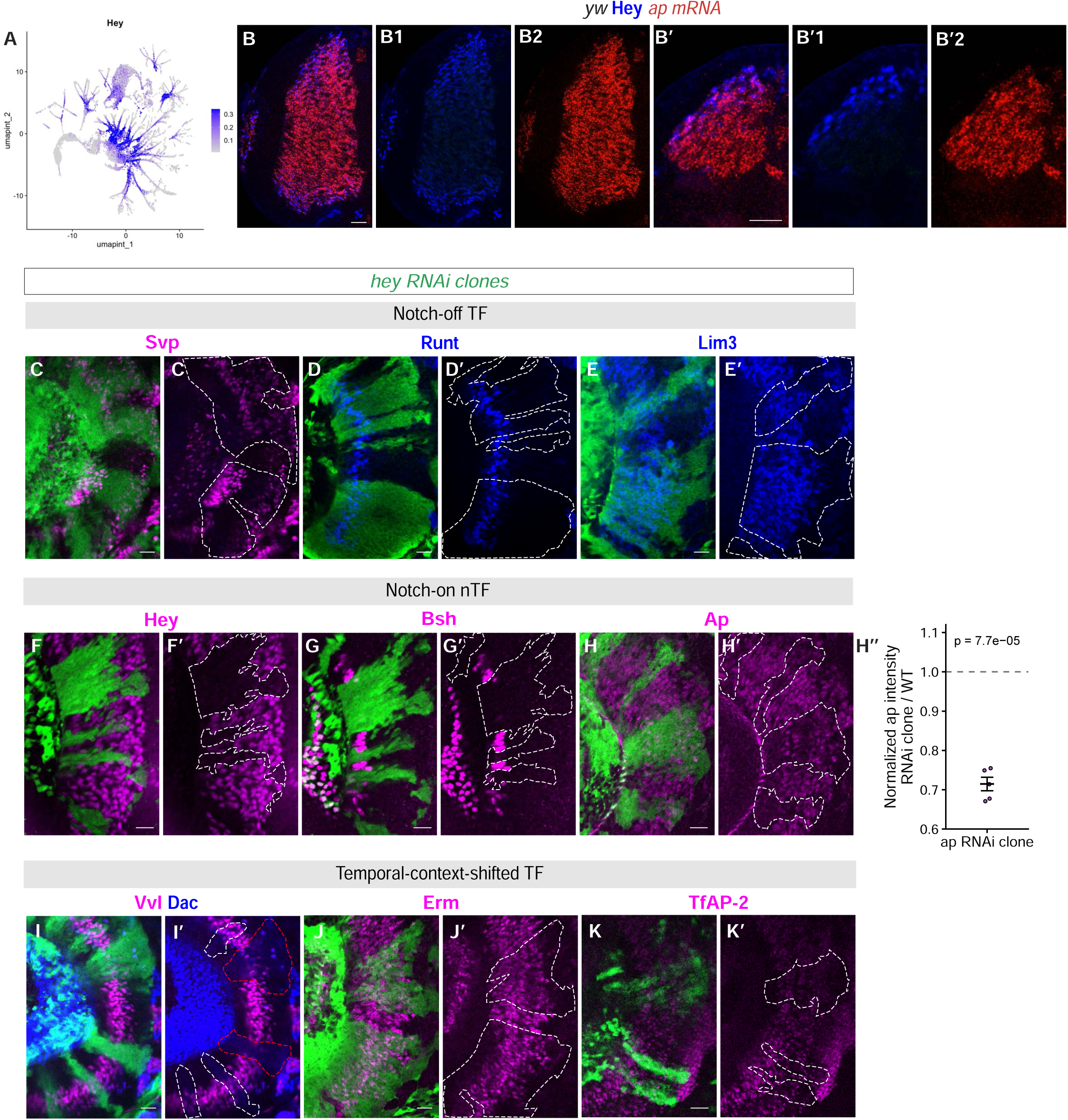
Hey is restricted to Notch-on immature neurons and controls only selected Notch-on nTF outputs. (A) Feature plot showing hey transcript expression in immature neuronal populations. (B) HCR RNA-FISH for ap mRNA combined with immunostaining for endogenous Hey protein in wild-type larval optic lobes. Hey protein is transiently detected in immature neurons and is restricted to ap+ Notch-on cells. Single-channel and magnified views are shown in B1–B′2. (C–E) Notch-off TF expression is not ectopically expanded in *hey* RNAi clones. Svp (C–C′), Run (D–D′), and Lim3 (E–E′) expression remain largely unchanged in clone regions. (F–H) Hey controls selected Notch-on nTF outputs. Hey expression is lost in *hey* RNAi clones (F–F′), Bsh expression is abolished (G–G′), and Ap expression is reduced but not eliminated (H–H′). (H″) Quantification of Ap intensity in *hey* RNAi clones. Ap intensity in RNAi clones was normalized to adjacent wild-type tissue from the same brain. Each point represents one brain with at least two clones measured. The dashed line indicates WT-normalized intensity = 1; mean ± SEM is shown. Significance was assessed by one-sample t test against 1 (p = 7.7 × 10⁻⁵). (I–K’) Temporal-context-shifted TFs show selective responses to hey depletion. Vvl expression is regionally reduced in regions marked by red dashed outlines but remains unchanged in regions marked by white dashed outlines (I–I′), whereas Erm (J–J′) and TfAP-2 (K–K′) are largely preserved. Dashed outlines mark the RNAi clone regions. Scale bars, 20 μm.

We next tested whether Hey broadly regulates Notch-on or Notch-off nTF expression. Because Hey is generally considered a transcriptional repressor, one possibility was that Hey represses Notch-off nTFs in Notch-on neurons. However, unlike *Su(H)* depletion, *Hey* RNAi did not cause ectopic expansion of Notch-off nTFs, including Svp, Run, Lim3, Ey, Hbn, and Oc (Figure 3C–E’ and Figure S2A–C’). Instead, loss of *Hey* affected selected Notch-on nTFs. In *Hey* RNAi clones, Bsh expression was abolished (Figure 3G-G’), whereas Ap were partially reduced (Figure 3H–H’). We found that Ap intensity was significantly reduced in *RNAi* clones compared with adjacent *wild type* (*wt*) tissue, with *RNAi* clones retaining 71.5 ± 1.7% of *wt* Ap signal (Figure 3H’’). Vvl showed regional loss within Notch-on cells (Figure 3I-I’). In contrast, Erm, TfAP-2, Scro, Toy, and Sox102F were largely unaffected (Figure 3J–3K’ and Figure S2D–F’).

Thus, contrary to the assumption, Hey does not repress Notch-off nTFs. Instead, Hey is required to activate the expression of selected Notch-on nTFs. However, Hey alone does not explain the broad requirement for Su(H) in Notch-on nTF expression. We therefore conclude that Hey functions as a selective Notch-on relay rather than a universal downstream mediator of canonical Notch signaling.

### An E-box-dependent enhancer explains the selective Hey requirement for *bsh*

We next asked how Hey controls *bsh*, the Notch-on nTF most strongly affected by *Hey* depletion. Bsh is expressed in and required for the specification of medulla Mi1 and two types of lamina neurons^31,36^. Using paired single-cell multiome data, we identified multiple differentially accessible elements linked to *bsh* expression, including *bsh*#2–#5 (Figure 4A). Feature plots of individual peak accessibility showed that these elements were accessible in *bsh*-expressing populations, including Mi1 and lamina neurons, supporting them as candidate regulatory elements (Figure 4B). Because these linked elements were associated with both medulla and lamina Bsh+ populations, we next used existing Janelia FlyLight GAL4 lines ^37,38^ to localize medulla-specific regulatory activity within the broader *bsh* locus.

**Figure 4.**
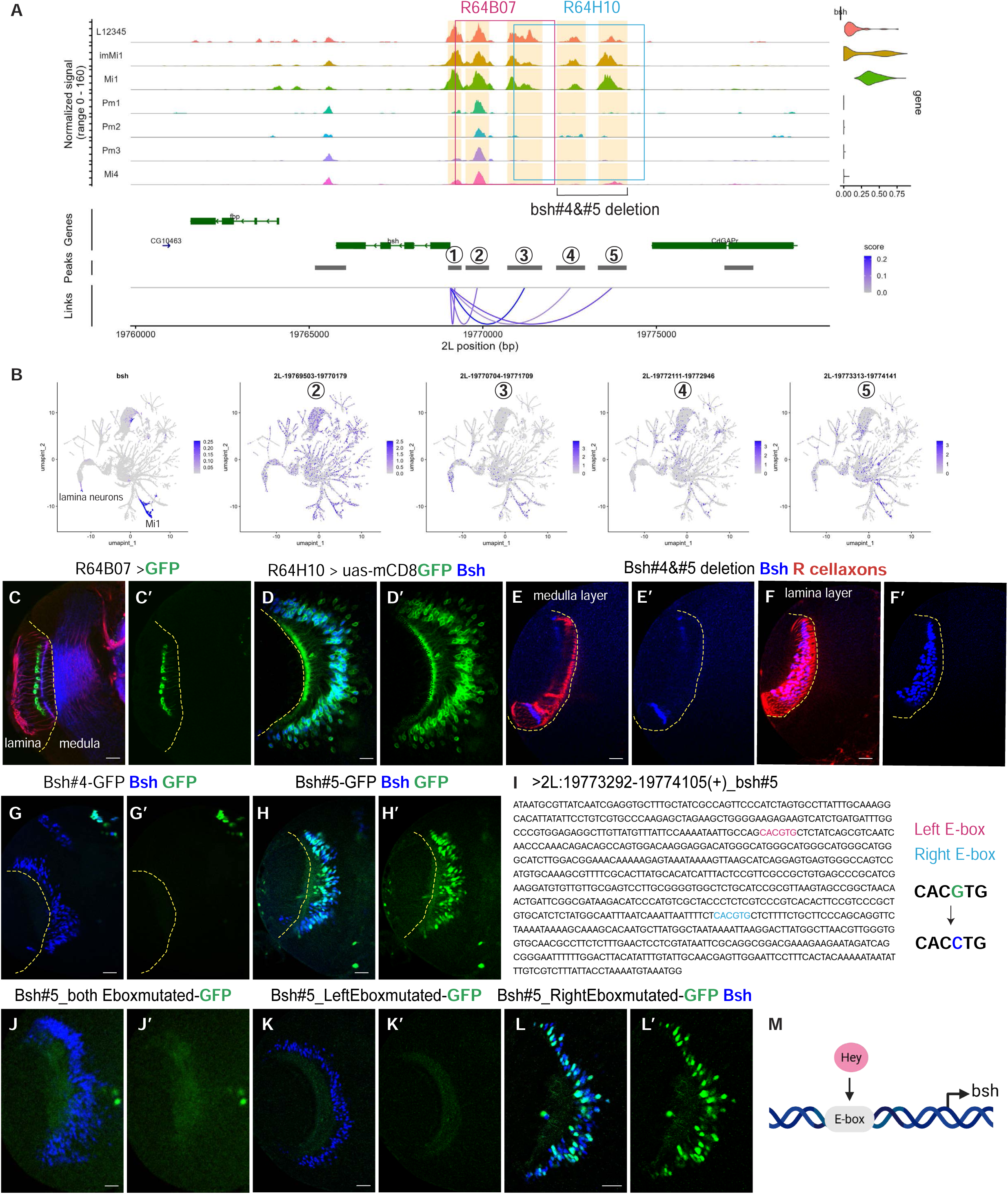
A Hey-sensitive E-box enhancer module controls medulla bsh expression. (A) Genome browser coverage plot showing chromatin accessibility across the bsh locus in optic lobe populations. Candidate accessible elements are numbered bsh#1–#5. Peak-gene links connect distal accessible elements to bsh expression. The genomic regions covered by Janelia FlyLight GAL4 lines R64B07 and R64H10 are indicated. (B) Feature plots showing bsh expression and accessibility of candidate linked peaks bsh#2–#5 on the scATAC UMAP. Candidate elements show accessibility in bsh-expressing populations, including Mi1 and lamina neurons. (C and D) Janelia GAL4 reporter analysis localizes medulla bsh regulatory activity to the downstream region. R64B07 drives reporter expression primarily in lamina Bsh+ cells but not in medulla Bsh+ Mi1 neurons (C–C′). In contrast, R64H10 drives reporter expression in medulla Bsh+ Mi1 neurons (D–D′). (E and F) Endogenous deletion of the bsh#4/#5 region abolishes medulla Bsh expression while preserving lamina Bsh expression. Medulla Bsh expression is lost in the deletion mutant (E–E′), whereas lamina Bsh expression remains detectable (F–F′). (G and H) Reporter assays testing individual candidate enhancers. bsh#4-GFP does not drive medulla Bsh+ expression (G–G′), whereas bsh#5-GFP is sufficient to drive reporter expression in medulla Bsh+ neurons (H–H′). (I) Sequence of the cloned bsh#5 enhancer fragment. Two Hey-class/E-box-like motifs are highlighted. The left E-box and right E-box motifs were mutated from CACGTG to CACCTG. (J–L’) E-box motif mutagenesis of bsh#5. Mutation of both E-box motifs abolishes reporter activity (J–J′). Mutation of the left E-box also abolishes reporter activity (K–K′), whereas mutation of the right E-box preserves reporter expression in Bsh+ medulla neurons (L–L′). (M) Model showing that Hey-dependent E-box input is required for bsh#5 enhancer activity and medulla bsh expression. Yellow dashed lines mark the boundary between the lamina and medulla regions. The lamina is positioned on the left and the medulla on the right. Scale bars, 20 μm.

The R64B07 GAL4 line, which covers the more upstream candidate region (Figure 4A), drove reporter expression in lamina Bsh+ cells but not in medulla Bsh+ Mi1 neurons (Figure 4C-C’). In contrast, the R64H10 GAL4 line, which covers the downstream region containing *bsh#4* and *bsh#5* (Figure 4A), drove strong reporter expression in medulla Bsh+ Mi1 neurons, with only sparse expression in lamina neurons (Figure 4D-D’). These results nominated the *bsh#4/#5* region as a candidate medulla bsh regulatory element.

Consistent with this, deletion of the endogenous *bsh#4/#5* region abolished medulla Bsh expression while preserving lamina Bsh expression, demonstrating that this region is specifically required for the medulla Bsh expression (Figure 4E-F’). We then tested the candidate elements individually in reporter assays. To minimize position effects, all reporter constructs were inserted into the same third-chromosome landing site. *bsh#5* was sufficient to drive medulla expression in the same pattern as endogenous Bsh (Figure 4H-H’), whereas *bsh#4* was not sufficient on its own (Figures 4G-G’). Thus, the *bsh#4/#5* region is required for medulla bsh expression, and *bsh#5* contains enhancer activity sufficient to drive this expression.

Motif analysis identified two Hey-class E-box motifs within *bsh#5* (Figure 4I). Mutation of both E-box motifs abolished the enhancer activity. Strikingly, despite the two motifs having identical sequences and each being altered by only a single central nucleotide (CACGTG to CACCTG), single-site mutagenesis revealed that one motif was uniquely required, as mutation of this site alone abolished reporter expression (Figures 4J–L’). Thus, medulla *bsh* expression is controlled by a dedicated enhancer module whose activity requires a specific E-box input.

Together with the *hey* RNAi results, these findings suggest that Hey directly activates *bsh* expression through an E-Box motif in *bsh-CRE#5*, which represents a Notch-on enhancer in which E-box input is essential (Figure 4M). Our motif analysis of *bsh* enhancer explains why it is particularly sensitive to Hey depletion. More broadly, these results support a model in which Hey dependency is determined not simply by motif enrichment across Notch-on chromatin, but by local enhancer architecture that converts E-box regulatory potential into essential enhancer activity.

### Sim/Tgo defines a parallel Notch-on relay with selective nTF output

Because Hey controlled only a subset of Notch-on nTFs, we next asked whether additional E-box-binding Notch-dependent relay factors control other Notch-on transcriptional outputs. Analysis of the RNA modality of the multiome data identified *sim*, encoding a bHLH-PAS transcription factor ^39^, as being expressed after the earliest Hey+ immature stage, in young differentiating Notch-on neurons that included Bsh^+^ Mi1 neurons, TfAP-2^+^ Notch-on Tm1, Tm2, Tm4, and Tm6 clusters, and Tm9 neurons (Figure 5A). Both a *sim-GFP* pBAC reporter and antibody staining for endogenous Sim showed the same expression pattern, confirming Sim expression in these Notch-on populations (Figure 5B-B’). By contrast, the RNA modality showed that Tgo, the obligate heterodimeric partner of Sim ^40^, was broadly expressed across medulla cell states (Figure 5A’). Thus, Sim likely provides the restricted spatial input for Sim/Tgo-dependent regulation, whereas Tgo is broadly available.

**Figure 5.**
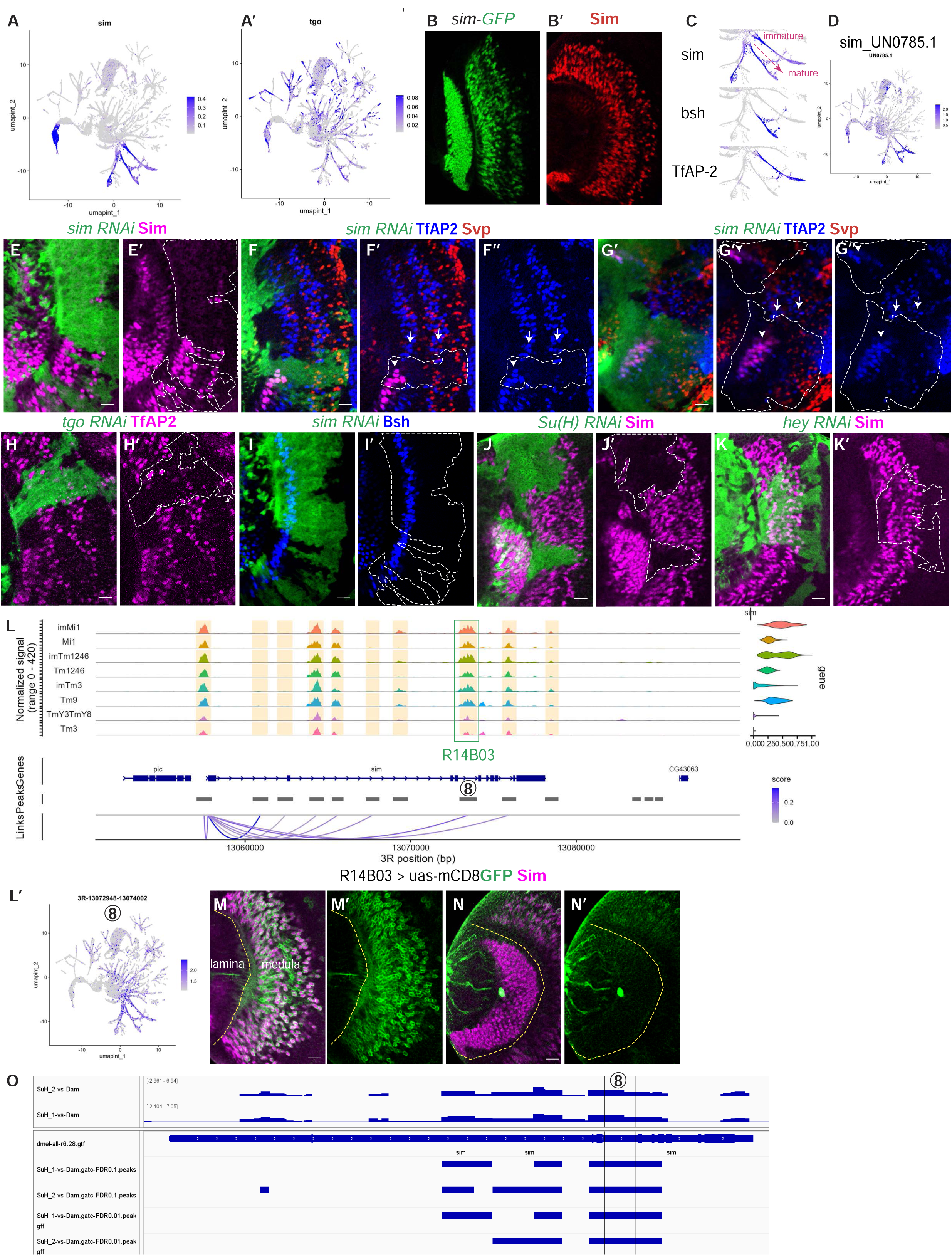
Sim/Tgo defines a parallel Notch-on bHLH relay that selectively controls TfAP-2. (A and A′) Feature plots showing expression of sim (A) and tgo (A′). sim is enriched in subsets of Notch-on populations, whereas tgo is broadly expressed across medulla cell clusters. (B and B′) Validation of Sim expression using a sim-GFP pBAC reporter and immunostaining for endogenous Sim protein. (C) Feature plots showing the relative timing of sim, bsh, and TfAP-2 expression along the neuronal differentiation trajectory. sim expression appears before robust bsh and TfAP-2 expression. (D) Feature plot showing chromVAR deviation scores for a Sim/Tgo-like motif, indicating enrichment of Sim/Tgo-associated motif activity in sim-expressing populations. (E–G) *sim* RNAi selectively disrupts Notch-on TfAP-2 expression. Sim expression is lost in *sim* RNAi clones (E–E′). In *sim* RNAi clones, TfAP-2 expression is lost in Notch-on cells, while Svp+ Notch-off cells are preserved (F–G′′). Arrows mark Notch-on TfAP-2+ cells and arrowhead marks Notch-off TfAP2+ cells. (H and H′) Depletion of tgo, the broadly expressed Sim heterodimeric partner, similarly abolishes TfAP-2 expression in clone regions. (I and I′) *sim* RNAi does not disrupt Bsh expression, indicating that Sim is not broadly required for all nTFs expressed in Sim+ neurons. (J and J′) Sim expression is lost in *Su(H)* RNAi clones, placing sim downstream of canonical Notch signaling. (K and K′) Sim expression is preserved in *hey* RNAi clones, indicating that sim is not downstream of Hey and instead defines a parallel Notch-dependent relay. (L) Genome browser coverage plot showing chromatin accessibility and peak-gene links at the sim locus. The candidate element sim#8 and the Janelia GAL4 fragment R14B03 spanning this region are indicated. (L′) Feature plot showing accessibility of sim#8. (M–N′) R14B03 drives reporter expression in Sim+ medulla cells but not in lamina cells. Yellow dashed lines mark the boundary between lamina and medulla regions. (O) Previously published Su(H) DamID tracks show Su(H) occupancy at the sim#8 region, supporting sim#8 as a candidate direct Su(H)-responsive enhancer. White dashed outlines mark RNAi clone regions. Scale bars, 20 μm.

The timing of sim expression suggested that Sim could act upstream of multiple Notch-on nTFs. Along the single-cell developmental trajectory, *sim* mRNA was detected before robust *bsh* and *TfAP-2* expression (Figure 5C). In addition, chromVAR analysis revealed elevated deviation scores for a Sim/Tgo type E-box-like motif in sim-expressing clusters, indicating that accessible elements in these cells are enriched for this E-box subtype (Figure 5D).

We first tested whether Sim is required for nTF expression in the Notch-on populations in which it is expressed. We found that *sim* RNAi strongly reduced TfAP-2 expression in Notch-on neurons (Figure 5F-G’’, arrow pointed stripe) while preserving TfAP-2 expression in Notch-off neurons (Figure 5F-G’’, arrowhead pointed stripe). Consistent with Sim acting through its broadly expressed obligate partner, depletion of *tgo* produced a similar loss of Notch-on TfAP-2 expression (Figure 5H-H’). In contrast, despite Sim expression in Mi1 neurons, loss of *sim* did not affect Bsh expression (Figure 5I-I’). In *sim* RNAi clones, *run*, which marks the Notch-off sibling of TfAP-2+ cells, was not ectopically detected in Notch-on cells (Figure S3A-A’), suggesting that loss of Sim does not transform TfAP-2+ Notch-on cells into their Notch-off siblings. Together, these results indicate that Sim/Tgo selectively controls the Notch-on component of TfAP-2, but not Bsh.

We next asked how Sim itself is activated downstream of Notch signaling. Multiome-based peak-gene linkage identified several candidate accessible elements associated with sim expression, including *sim#8* (Figure 5L-L’). A Janelia GAL4 line spanning *sim#8* drove reporter expression in Sim+ cells in the medulla (Figure 5M–M′), but not in the lamina (Figure 5N–N′), suggesting that *sim#8* contains regulatory activity specific to medulla Sim+ neurons. Moreover, previously published Su(H) DamID data ^41^ showed Su(H) occupancy at this enhancer region, supporting *sim#8* as a candidate direct Su(H)-responsive enhancer (Figure 5O). Consistent with this model, Sim expression was abolished in *Su(H)* RNAi clones (Figure 5J-J’). In contrast, Sim expression was not affected by *Hey* RNAi, indicating that Sim is not downstream of Hey but instead defines a parallel Notch-dependent relay (Figure 5K-K’).

Together, these results identify Sim/Tgo as a Notch-dependent E-box-binding relay that acts in parallel to Hey. Although Sim is expressed in multiple Notch-on populations and Sim/Tgo-like E-box motif accessibility is enriched in these cells, Sim/Tgo selectively controls the Notch-on component of TfAP-2 expression and is dispensable for Bsh. Thus, as with Hey, relay-factor expression and motif enrichment do not predict universal dependency. Instead, Notch-on nTF outputs are decoded selectively: Hey is required for Bsh, whereas Sim/Tgo is required for the Notch-on expression of TfAP-2.

### A transient *TfAP-2* enhancer contains separable Notch-on and Notch-off regulatory logic

We next asked how the Sim/Tgo relay is interpreted at the enhancer level. In previous work, we identified a Janelia GAL4 R24B02 fragment spanning the *TfAP-2#3* region that drove reporter expression in TfAP-2^+^ cells ^42^. Inspection of this region in our scATAC-seq data revealed that *TfAP-2#3* was the only accessible peak within the R24B02 fragment, making it the strongest candidate enhancer underlying the reporter activity (Figure 6A). Although *TfAP-2#3* was not linked to *TfAP-2* by multiome-based peak-to-gene analysis (Figure 6A), feature plots showed that its accessibility was specifically enriched in young TfAP-2^+^ neurons and was not maintained at later stages (Figure 6B), even though endogenous TfAP-2 expression persisted (Figure 6B’). This temporal accessibility pattern suggested that *TfAP-2#3* may act as an early enhancer for *TfAP-2* expression. We therefore cloned *TfAP-2#3* to test whether this transiently accessible element was sufficient to drive reporter expression in TfAP-2^+^ neurons.

**Figure 6.**
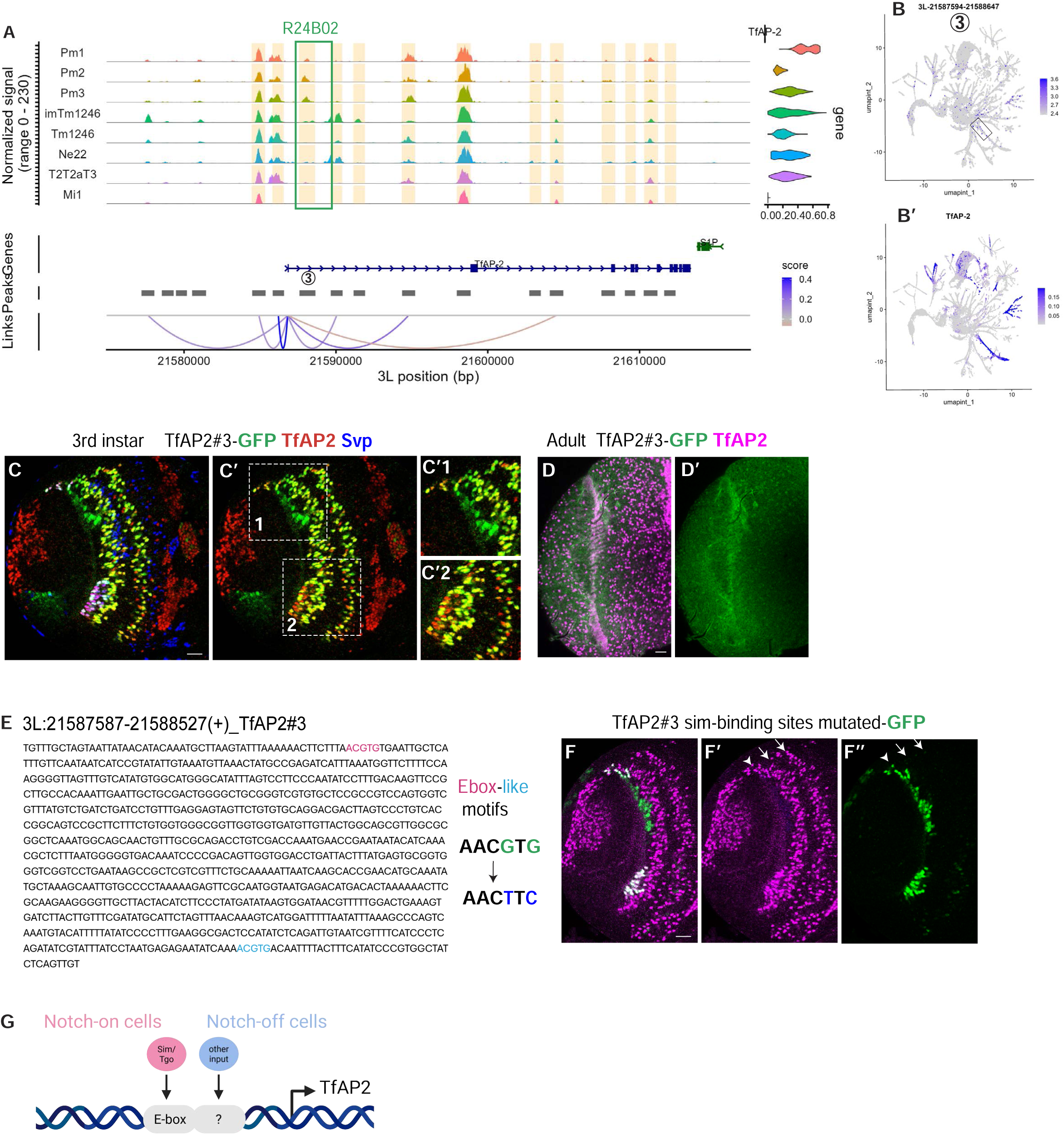
A transient TfAP-2 enhancer contains separable Notch-on and Notch-off regulatory logic. (A) Genome browser coverage plot showing chromatin accessibility and peak-gene links at the TfAP-2 locus. The previously identified Janelia GAL4 fragment R24B02 spans the TfAP-2#3 peak region. (B and B′) Feature plots showing accessibility of TfAP-2#3 (B) and TfAP-2 expression (B′). TfAP-2#3 is accessible in young TfAP-2+ neuronal populations, indicated by the boxed region. (C–C″) TfAP-2#3-GFP reporter expression in third-instar larval optic lobes. TfAP-2#3 drives reporter expression in major larval TfAP-2+ domains, including both TfAP-2+ Svp− Notch-on cells and TfAP-2+ Svp+ Notch-off cells. Magnified views of boxed regions are shown in C″1 and C″2. (D and D′) TfAP-2#3-GFP reporter expression in adult optic lobes. Although endogenous TfAP-2 expression persists in the adult, the TfAP-2#3 reporter is not maintained, indicating that TfAP-2#3 functions as a transient developmental enhancer. (E) Sequence of the cloned TfAP-2#3 enhancer fragment. Two Sim/Tgo-like E-box motifs are highlighted, and the motif mutations are indicated. (F–F″) Mutation of Sim/Tgo-like motifs in TfAP-2#3 selectively disrupts reporter activity in Notch-on TfAP-2+ cells while preserving reporter activity in Notch-off cells. Arrows indicate Notch-on TfAP-2+ cells lacking reporter expression after motif mutation and arrowhead indicate Notch-off cells with preserved reporter activity. (G) Model showing that TfAP-2#3 contains separable Notch-on and Notch-off regulatory inputs. Sim/Tgo-like E-box motifs are required for the Notch-on component, whereas the Notch-off component is controlled by an independent input. Scale bars, 20 μm.

*TfAP-2#3* was sufficient to drive reporter expression in major larval TfAP-2^+^ domains, including both Notch-on (Svp^-^) cells born in the two Opa temporal stages and Notch-off (Svp^+^) cells born in the Hth temporal stage ^42^ (Figure 6C). However, *TfAP-2#3* did not label every Notch-off (TfAP-2^+^ Svp^+^) cell (Figure 6C’2), and reporter activity was also observed in some TfAP-2^-^ cells likely born at the same temporal stage (Figure 6C’1). This pattern suggests that *TfAP-2#3* captures regulatory information associated with a broader temporal domain and Notch status, while for TfAP-2 expression in Notch-off cells, additional inputs are required to fully restrict the expression pattern. Consistent with the transient opening of this CRE, the *TfAP-2#3* reporter failed to maintain adult expression, despite persistent endogenous TfAP-2 expression (Figure 6D,D’). These results indicate that *TfAP-2#3* functions as a transient developmental enhancer and suggest that additional regulatory elements are required to refine and sustain TfAP-2 expression during later stages. We next asked whether the Notch-on and Notch-off components of *TfAP-2#3* activity are encoded by separable motif inputs. Within *TfAP-2#3*, we identified two Sim/Tgo-type E-box motifs (Figure 6E). Mutation of these motifs selectively abolished reporter activity in Notch-on TfAP-2^+^ cells (Figure 6F–F″, arrows-pointed stripes), while preserving reporter activity in Notch-off cells (Figure 6F–F″, arrowheads-pointed stripe). This selective disruption of reporter activity in Notch-on cells phenocopies the requirement for Sim/Tgo in endogenous Notch-on TfAP-2 expression (Figure 6F). Thus, the Notch-on and Notch-off components of *TfAP-2#3* activity are controlled by separable cis-regulatory inputs within the same enhancer.

These results extend the relay model to the enhancer level. Sim/Tgo is required for the Notch-on component of TfAP-2 expression, and Sim/Tgo-type E-box motifs within *TfAP-2#3* are specifically required for the reporter activity in Notch-on cells. In contrast, preservation of the Notch-off reporter component after Sim/Tgo motif mutation shows that the Notch-off component is encoded independently (Figure 6G). Thus, *TfAP-2#3* separates developmental timing from Notch-status logic: it acts transiently during larval development, but within that window it integrates separable cis-regulatory inputs for Notch-on and Notch-off expression.

### Tap selectively controls early-born Notch-off tup expression

We next asked whether relay specificity also operates in Notch-off neurons. Analysis of the single-cell RNA profiles showed that *tap*, encoding another bHLH family member, was broadly expressed in GMCs but was maintained only in immature early-born Notch-off neurons (Figure 7A). This pattern suggests that *tap* transcription is actively restricted during the transition from GMCs to neuronal progeny: it is turned off in Notch-on progeny and is not maintained in late-born Notch-off neurons. We validated *tap* expression using HCR FISH; consistent with the single-cell RNA data, *tap* was transiently transcribed (Figure 7E-F’) and its signal was lost in *tap* RNAi clones (Figure 7G-G’).

**Figure 7.**
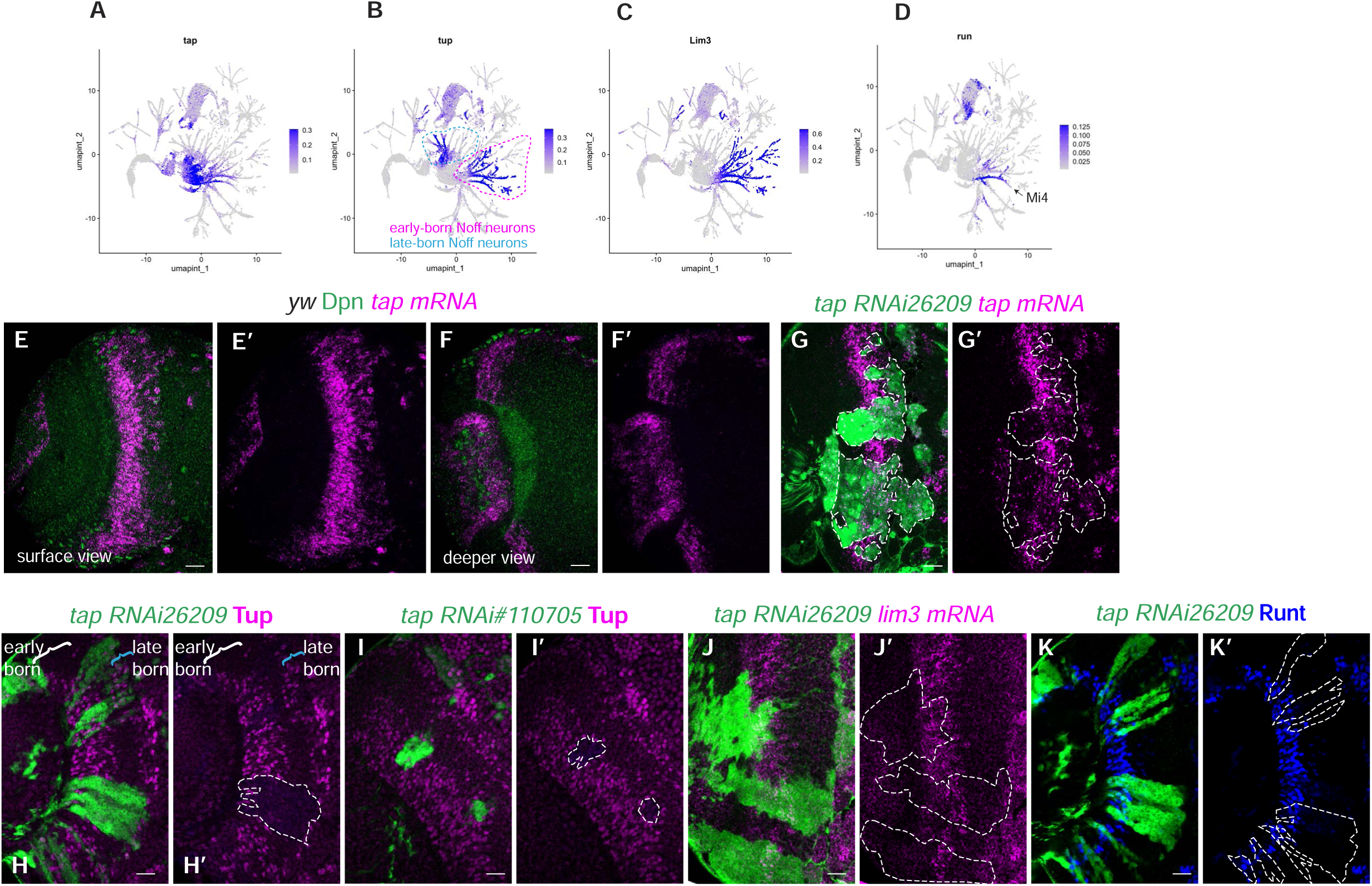
Tap selectively controls early-born Notch-off tup expression. (A–D) Feature plots showing expression of tap (A), tup (B), Lim3 (C), and run (D). tap is enriched in early-born Notch-off neurons. tup is expressed in both early-born and late-born Notch-off populations, whereas Lim3 is broadly expressed in early-born Notch-off neurons and run is restricted to Mi4. (E–F′) HCR RNA-FISH for tap mRNA in wild-type larval optic lobes. tap transcripts are detected transiently in GMCs and immature neurons, shown in surface and deeper optical sections. Dpn labels neuroblasts. (G and G′) tap mRNA is reduced in tap RNAi clones, confirming RNAi-mediated depletion. (H–I′) *tap* RNAi selectively disrupts early-born tup expression. In *tap* RNAi clones, early-born Tup expression is lost, whereas late-born Tup expression is preserved. Two independent tap RNAi lines are shown. (J and J′) Lim3 expression in early-born Notch-off neurons is preserved in *tap* RNAi clones. (K and K′) Run expression is preserved in *tap* RNAi clones. White dashed outlines mark RNAi clone regions. Scale bars, 20 μm.

Early-born Notch-off neurons include multiple neuron types, such as Pm neurons, Mi4, Mi9, and Dm10. To test whether Tap broadly regulates early-born Notch-off identity or instead controls selective transcriptional outputs, we examined several nTFs expressed in this population. Tup (Figure 7B) and Lim3 (Figure 7C) are broadly expressed in early-born Notch-off neurons, whereas run is restricted to Mi4 (Figure 7D). Tup is also expressed in an additional late-born Notch-off domain (Figure 7B).

In *tap* RNAi clones, early-born tup expression was selectively lost, whereas late-born tup expression remained intact (Figure 7H-I’). In contrast, Lim3 expression in early-born Notch-off neurons was unaffected (Figure 7J-J’). Run, which specifically marks Mi4, was also preserved after *tap* RNAi (Figure 7K-K’). Thus, Tap is not required for early-born Notch-off identity as a whole. Instead, Tap selectively controls early-born Tup expression.

These results show that relay specificity is not unique to the Notch-on state. Even within early-born Notch-off neurons, Tap selectively controls one nTF output, *tup*, while leaving other early Notch-off markers intact. Thus, both Notch-on and Notch-off programs are organized into selective relay-dependent modules.

### Distinct *tup* enhancers separately control early- and late-born Notch-off expression

Because loss of Tap selectively affected early-born Tup expression without disrupting the late-born Tup domain, we asked whether these two expression domains are regulated by distinct enhancer modules. Single-cell chromatin accessibility analysis identified candidate regulatory elements near the *tup* locus with different temporal accessibility patterns (Figure 8A). *tup#14* was preferentially accessible in early-born Tup^+^ Notch-off neurons, whereas *tup#20* was preferentially accessible in late-born Tup^+^ Notch-off neurons (Figure 8A-B). Consistent with the late accessibility of *tup#20*, the Janelia GAL4 line R76B03, which spans the *tup#20* region, drove reporter expression in late-born Tup^+^ cells (Figure 8C).

**Figure 8.**
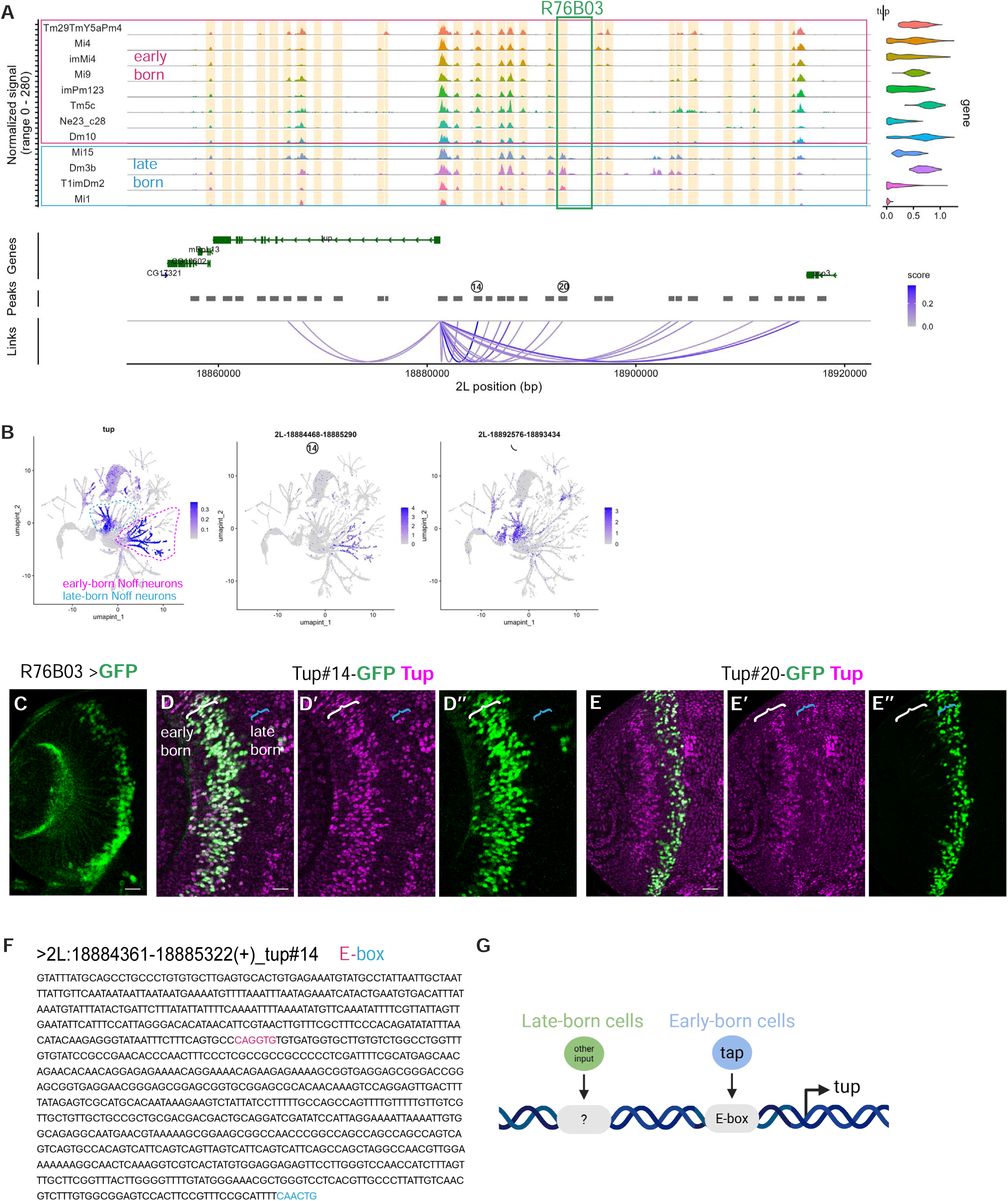
Distinct tup enhancers separate early- and late-born Notch-off expression. (A) Genome browser coverage plot showing chromatin accessibility and peak-gene links at the tup locus across early-born and late-born Notch-off neuronal populations. Candidate elements tup#14 and tup#20 are indicated. The Janelia GAL4 fragment R76B03, which spans the tup#20 region, is highlighted. (B) Feature plots showing tup expression and accessibility of candidate elements tup#14 and tup#20. tup#14 is preferentially accessible in early-born tup+ Notch-off neurons, whereas tup#20 is preferentially accessible in late-born tup+ Notch-off neurons. (C) R76B03 > GFP reporter expression, showing activity consistent with the late-born tup expression domain. (D–D″) tup#14-GFP reporter expression in larval optic lobes. tup#14 drives reporter expression preferentially in early-born Tup+ Notch-off neurons. (E–E″) tup#20-GFP reporter expression in larval optic lobes. tup#20 drives reporter expression preferentially in late-born Tup+ Notch-off neurons. (F) Sequence of the cloned tup#14 enhancer fragment. Predicted E-box-like motifs are highlighted. (G) Model showing that early- and late-born tup expression domains are encoded by distinct enhancer modules. The early-born domain is driven by tup#14 and is Tap-dependent, whereas the late-born domain is driven by tup#20 and is regulated independently of Tap. White brackets indicate early-born regions, and blue brackets indicate late-born regions. Scale bars, 20 μm.

We next tested these candidate elements directly in reporter assays. *tup#14* drove reporter expression in early-born Tup^+^ Notch-off neurons (Figures 8D-8D’’), whereas *tup#20* drove expression preferentially in late-born Tup^+^ Notch-off neurons (Figures 8E-E’’). Thus, early and late tup expression domains are not generated by a single shared enhancer. Instead, temporally separable enhancer modules independently encode distinct components of *tup* expression. Notably, we identified two E-box-like motifs in the *tup#14* enhancer, which differ from the Hey type E-box in the two central bases, and could be potentially bound by Tap according to the Tap consensus motif, providing candidate inputs for future functional testing (Figure 8F).

These findings provide a cis-regulatory explanation for why Tap controls early-born *tup* expression without affecting late-born *tup* expression: the two expression domains are encoded by distinct enhancer modules. More broadly, these results support a model in which selective nTF dependency reflects modular enhancer organization, with different expression domains of the same nTF encoded by distinct regulatory elements and potentially distinct upstream inputs (Figure 8G).

### *Lim3* expression is reconstructed by combining temporal and Notch-status-specific enhancer inputs

We next asked whether temporal information and Notch status can also be encoded by separate enhancer modules that act together to generate a precise nTF expression pattern. *Lim3* provided a useful test case because it is expressed in Notch-off neurons within a defined temporal window.

The Janelia GAL4 line 33H04 near the *Lim3* locus drove reporter expression broadly within the corresponding temporal domain but lacked Notch-off specificity (Figure 9A and 9D,D’). Peak accessibility analysis of the 33H04-covered region (*#9* and *#10*) supported this element as a temporal identity regulatory candidate since it opens in both notch-on and notch-off progenies born at the early-temporal stages (Figure 9B-9B’’). We therefore searched the scATAC-seq data for candidate Notch-off-associated regulatory elements near *Lim3*. We selected a candidate Notch-off element (*#13*) because it was preferentially accessible in Lim3+ Notch-off neurons, was positioned near the *Lim3* locus, and contained E-box-like motifs, that differ from Hey type E-Box in the two central bases, consistent with the bHLH relay theme (Figure 9A, 9C and 9C’).

**Figure 9.**
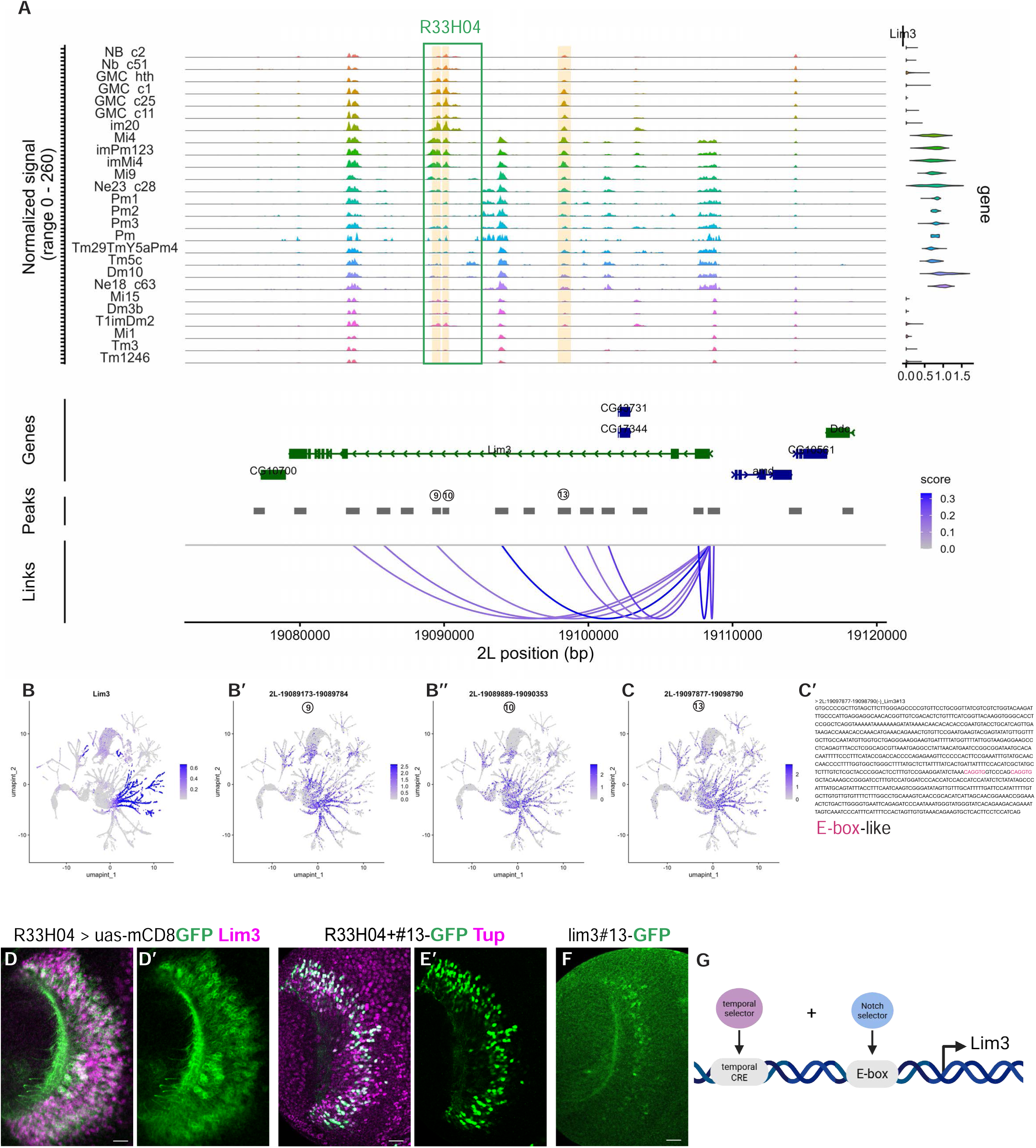
Lim3 expression is reconstructed by combining temporal and Notch-status-specific enhancer inputs. (A) Genome browser coverage plot showing chromatin accessibility and peak-gene links at the lim3 locus across developing medulla cell states. The Janelia GAL4 fragment R33H04 and candidate linked elements lim3#9, lim3#10, and lim3#13 are indicated. (B–C) Feature plots showing lim3 expression (B) and accessibility of candidate linked elements lim3#9 (B′), lim3#10 (B″), and lim3#13 (C). Lim3#13 is preferentially accessible in Lim3+ Notch-off neurons. (C′) Sequence of the cloned lim3#13 element. Predicted E-box-like motifs are highlighted. (D and D′) R33H04 drives reporter expression broadly within the Lim3 temporal domain, including cells that do not express endogenous Lim3, suggesting that this element primarily encodes temporal regulatory information but lacks full Notch-off specificity. (E and E′) Combining R33H04 with lim3#13 drives reporter expression that more closely matches endogenous Lim3 expression in Notch-off neurons. Tup has exactly the same expression pattern in early-born Notch-off neurons as Lim3, and is used here because Lim3 antibody has run out. (F) Lim3#13-GFP alone shows weak or limited reporter activity, indicating that the Notch-off-associated element is not sufficient on its own to fully drive the endogenous Lim3 pattern. (G) Model showing that precise lim3 expression is generated by combinatorial enhancer integration: the R33H04 element provides temporal information, whereas lim3#13 contributes Notch-off-associated regulatory input. Scale bars, 20 μm.

Combining the 33H04 temporal element with the candidate Notch-off-associated element (*#13*) produced reporter activity that more closely recapitulated endogenous Lim3 expression (Figure 9E–E′). These results suggest that precise Lim3 expression is generated by integrating distinct enhancer inputs, with one element providing temporal information and the other contributing Notch-off specificity. Notably, the candidate Notch-off-associated element (*#13*) alone showed only weak enhancer activity (Figure 9F), indicating that its activity may depend on cooperation with the temporal element. The presence of E-box-like motifs, that differ from the Notch-on Hey type E box, within the Notch-off-associated element suggests that E-box-dependent regulatory logic may also contribute to Lim3 restriction (Figure 9G).

Interestingly, this regulatory principle may be used by additional genes. *CG11294* exhibits an expression pattern similar to *Lim3*, and an enhancer encompassed by the Janelia GAL4 line R83D05 drives activity in both Notch-on and Notch-off cells born within the same temporal window (Figure S4A-C’). This suggests that *CG11294* may also rely on a combinatorial enhancer strategy, in which temporal information is provided by one element and Notch-off specificity is imposed by another. In support of this model, we identified a highly linked candidate CRE (*#5*) as a potential Notch-off element. Notably, this element contains multiple E-box-like motifs that differ from the Hey type E box in the two central bases (Figure S4A,B,D).Together, these findings expand the modular enhancer model beyond single-enhancer logic. While *bsh#5* and *TfAP-2#3* illustrate how individual enhancers can encode specific relay or Notch-status inputs, *Lim3* demonstrates that these regulatory inputs can also be distributed across separate cis-regulatory elements and assembled combinatorially to generate precise neuron-type-specific expression.

## DISCUSSION

A central challenge in developmental biology is to understand how a limited set of signaling pathways generate diverse developmental outcomes. Here, we show that Notch signaling diversifies medulla neurons through modular relay-enhancer decoding, rather than through a single linear downstream pathway. Canonical Notch/Su(H) provides a broad Notch-status input, but individual neuron-type transcription factor outputs are selected by distinct E-box-associated bHLH family relay factors and local enhancer architectures (Figure 10). This organization allows a binary Notch-on/Notch-off sibling fate decision to be interpreted in temporal and spatial contexts, thereby generating diverse nTF programs during medulla development.

**Figure 10.**
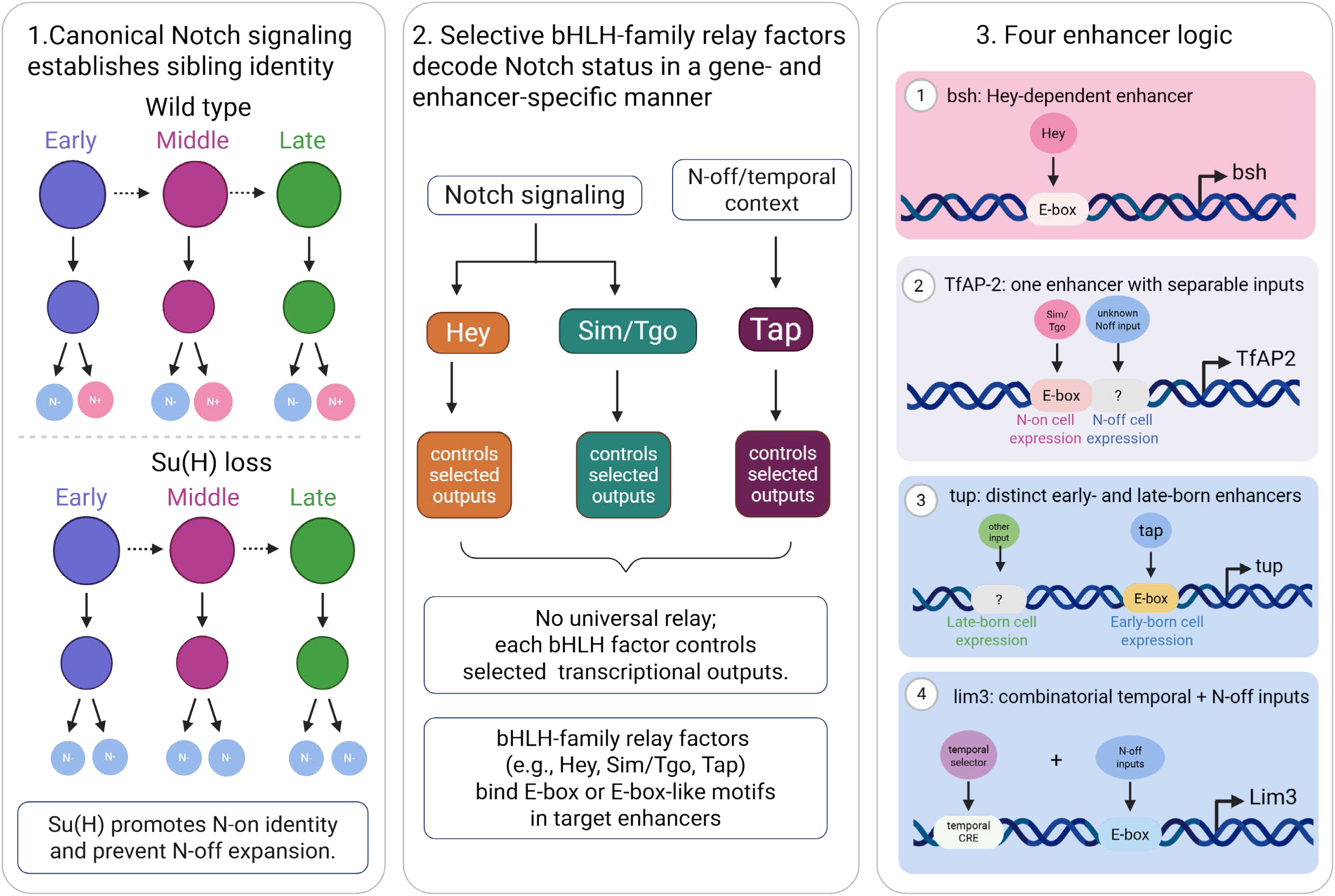
Selective bHLH relay factors and modular enhancers decode temporal and Notch-status inputs into nTF outputs. Left: Model of canonical Notch/Su(H) function across neuroblast temporal stages. In wild type, neuroblasts progress through early, middle, and late temporal windows, and each GMC division generates a Notch-off sibling and a Notch-on sibling. These two inputs—temporal stage and Notch status—produce distinct nTF programs in each temporal window. Upon loss of Su(H), Notch-on nTF programs are lost and Notch-off nTF programs expand, indicating that canonical Notch signaling promotes Notch-on identity and prevents inappropriate Notch-off expansion. Middle: Notch-status information is decoded by selective bHLH-family relay factors rather than by a single universal downstream effector. Hey and Sim/Tgo act in Notch-on contexts, whereas Tap acts in an early-born Notch-off/temporal context. Each relay factor controls selected transcriptional outputs in a gene- and enhancer-specific manner. Right: Four regulatory principles illustrate how enhancer architecture determines relay specificity. Bsh is controlled by a Hey-dependent E-box enhancer. TfAP-2 contains separable Notch-on and Notch-off enhancer inputs, with Sim/Tgo required for the Notch-on component. tup expression is separated into early- and late-born enhancer modules, with Tap controlling the early-born component. Lim3 expression is generated by combining temporal and Notch-off-associated enhancer inputs. Together, the model proposes that medulla neuronal diversity is generated by the intersection of neuroblast temporal stage and Notch status, which is decoded through selective bHLH relay factors and modular cis-regulatory elements.

The Hey and Hes family Notch target genes are generally known to encode transcriptional repressors ^17,19,20^. Since loss of Notch signaling results in loss of Notch-on nTF expression and ectopic Notch-off nTF expression, the initial hypothesis was that Hey should repress Notch-off gene expression in Notch-on neurons. However, to our surprise, Hey is not required to repress Notch-off genes but required to activate the expression of Notch-on nTFs. Our findings also revise a simple model in which Su(H) activates one common Notch-on effector, such as Hey, to control a uniform Notch-on transcriptional program. Although Su(H) is broadly required for Notch-on nTF expression and prevents ectopic Notch-off programs, individual outputs are controlled by different relay factors. Hey is required for *bsh*, Sim/Tgo is required for the Notch-on component of *TfAP-2*, and Tap selectively controls early-born *tup* expression. Thus, Notch-dependent neuronal diversification is achieved by distributing Notch-status information across relay factors that act in specific temporal and spatial contexts.

A striking feature of these relay factors is that Hey, Sim, and Tap are all bHLH or bHLH-related transcription factors associated with E-box-type regulatory logic. These findings together suggest that Notch-dependent neuronal specification is mediated, at least in part, through a diversified bHLH/E-box relay layer. Hey functions in immature Notch-on neurons and is essential for the “CACGTG” type E-box-dependent *bsh#5* enhancer activity. Sim, together with broadly expressed Tgo, defines a parallel Notch-on relay that controls the Notch-on component of TfAP-2 through a variant E box”AACGTG” in *TfAP-2 #3*. Tap extends relay specificity to early-born Notch-off neurons by selectively controlling early tup expression. Notably, the E-Boxes present in the enhancers of Notch-off nTF genes that we cloned are never the Hey type E-box, but differ in the two central nucleotides. Interestingly, a recent study in the *Drosophila* ventral nerve cord identified a bHLH factor, Fer3, which binds a non-Hey type E-Box, and, and functions as an activator to promote Notch-off identity ^43^. Taken together, these findings suggest related bHLH/E-box regulatory mechanisms can be reused and diversified to generate distinct nTF outputs downstream of a common Notch-status input.

Our scATAC-seq demonstrated dramatic differences in the accessible chromatin landscapes in Notch-on vs Notch-off hemilineages. These results also clarify how motif enrichment should be interpreted. Hey type E-box motif accessibility is broadly associated with Notch-on chromatin, yet only selected Notch-on nTFs require Hey, suggesting possible redundancy between multiple bHLH factors that bind to the Hey type E-Box. The obvious candidates include the *E(spl)* family genes. However, due to the great number of *E(spl)* family genes and the difficulty to remove all of them, we were not able to determine if they function redundantly with Hey. Sim is expressed in multiple Notch-on neuron types, but Sim/Tgo is required for TfAP-2 and not Bsh. Thus, motif accessibility marks regulatory potential, whereas functional dependency is determined by enhancer architecture and trans-factor environment.

Our results demonstrate that enhancer modularity provides the mechanism for regulatory specificity. At the *bsh* locus, *bsh#5* contains Hey type E-box which is critical for enhancer activity, explaining why Bsh is highly sensitive to Hey depletion. At the *TfAP-2* locus, the transient *TfAP-2#3* enhancer drives both Notch-on and Notch-off larval expression, but Sim/Tgo-like motif mutation selectively removes the Notch-on component. At the *tup* locus, distinct enhancers encode early- and late-born expression, explaining why Tap affects early Tup without disrupting the late Tup domain. For Lim3, temporal and Notch-off information are distributed across separate elements that together better reconstruct endogenous expression. Together, these examples show that enhancers act as decoding units that integrate Notch status, temporal context, and relay-factor input to generate precise nTF expression.

Several questions remain. First, chromVAR and motif analyses identify candidate regulatory inputs, but they do not prove direct binding or functional requirement. Cell-type-specific CUT&RUN, CUT&Tag, or DamID will be needed to define where Hey, Sim/Tgo, and Tap bind *in vivo*. Second, although *sim#8* is supported by peak-gene linkage, reporter activity, Su(H) DamID occupancy, and loss of Sim after *Su(H)* depletion, mutating Su(H)-responsive sites or deleting *sim#8* would be needed to prove direct Su(H)-dependent enhancer regulation. Third, the Tap–tup module remains less mechanistically resolved than the Hey–bsh and Sim/Tgo–TfAP-2 modules. Testing whether *tup#14* reporter activity depends on Tap, or whether Tap-associated motifs are required, would clarify whether Tap directly controls the early *tup* enhancer.

Finally, future work should test whether this relay-enhancer logic applies genome-wide. A key question is whether other Notch-on and Notch-off nTFs are controlled by additional relay factors that recognize distinct motif classes enriched in status-associated open chromatin. Although Hey, Sim/Tgo, and Tap each regulate only a limited set of nTF outputs, this selectivity likely reflects a broader organizational principle rather than an exception. The Notch-on chromatin landscapes contain many accessible elements enriched for E-box or E-box-like motifs, suggesting that additional bHLH-family factors or related transcriptional cofactors may act as relay factors for other nTFs. In this view, Hey, Sim/Tgo, and Tap represent examples of a larger relay network in which different motif subtypes, temporal windows, and enhancer architectures specify which nTFs respond in each cellular context. Systematic comparison of motif enrichment in Notch-on and Notch-off accessible regions, combined with chromVAR activity, peak-gene linkage, enhancer perturbation, and relay-factor knockdown, could reveal whether each nTF output is associated with a specific relay-motif-enhancer grammar. Such analyses may also distinguish enhancers that are merely made accessible from those that are functionally dependent on a particular relay factor. This would extend the current gene-by-gene examples into a global regulatory model for how temporal stage and Notch status are decoded across the medulla. Together, our study shows that Notch-dependent nTF outputs are specified through selective relay factors and modular enhancer architecture, providing a mechanism by which a binary fate signal generates diverse neuronal identities.

Interestingly, Notch signaling also plays central roles in vertebrate neural development, where it regulates progenitor maintenance, neuronal differentiation, and cell-fate specification in a highly context-dependent manner^44,45^. Moreover, vertebrate Notch signaling has been shown to engage distinct transcriptional programs in different cellular environments through interactions with lineage-specific cofactors and chromatin landscapes^46^. These observations raise the possibility that the principles uncovered here may extend beyond the fly visual system.

More broadly, our findings suggest a general mechanism for developmental signaling specificity. Reused signaling pathways such as Notch, Wnt, Hedgehog, BMP, and EGFR must generate different outputs across tissues and developmental stages. Our results support a model in which specificity is not determined simply by pathway activity itself, but by how that activity is interpreted by relay factors and enhancer architecture in each cellular context. In this framework, a common signal provides a broad input, relay factors refine that input, and enhancers determine which target genes respond. The *Drosophila* medulla therefore provides a tractable example of how a shared developmental signal can generate diverse transcriptional outputs through modular relay-enhancer decoding. We propose that developmental signaling pathways may commonly achieve specificity through a hierarchical regulatory framework in which signaling effectors act through intermediate relay factors, and enhancer architecture integrates these inputs to generate context-dependent gene expression programs. Whether such relay-enhancer decoding mechanisms operate in vertebrate systems will be an important question for future studies.

## METHODS

### Fly stocks and husbandry

Flies were maintained on standard cornmeal medium under standard laboratory conditions. Unless otherwise indicated, crosses were maintained at 25°C. The following RNAi lines were used: UAS-Su(H) RNAi (VDRC 103597), UAS-Hey RNAi (BDSC 31891), UAS-sim RNAi (BDSC 26739), UAS-tgo RNAi (BDSC 38928), and UAS-tap RNAi (BDSC 26209). GFP-tagged BAC lines included hey-GFP (BDSC 64826) and sim-GFP (BDSC 93086). Janelia FlyLight GAL4 lines used in this study included R64B07 (BDSC 39293), R64H10 (BDSC 39326), R14B03 (BDSC 49254), R24B02 (BDSC 49312), R76B03 (BDSC 47751), and R33H04 (BDSC 48118).

### Generation of mosaic RNAi clones

Mosaic RNAi clones were generated using the ayGal4 FLP-out system. Virgin females of genotype yw hs-FLP; act>y+>Gal4 UAS-GFP/CyO; UAS-Dcr2/TM6B were crossed to males carrying individual UAS-RNAi transgenes. Progeny were raised at 25°C and heat shocked once at 37°C for 8 min during the first-instar larval stage to induce FLP-mediated excision of the FRT-flanked STOP cassette and activate actin-Gal4-driven RNAi expression. Larvae were then transferred to 29°C for 3 days before dissection at the wandering third-instar stage.

### Preparation of optic lobe cells for single-cell multiome and single-nucleus ATAC-seq

For single-cell multiome experiments, approximately 120 GFP-positive third-instar larvae were collected. Larvae were washed twice with PBS, sterilized briefly in 70% ethanol for 1 min, and rinsed again with PBS. Brains were dissected on ice in complete Schneider’s medium consisting of Schneider’s insect medium supplemented with 10% fetal bovine serum, 2% penicillin-streptomycin, and 0.02 mg/mL insulin. Eye discs were removed during dissection. Dissected brains were transferred to a glass dish containing Dulbecco’s phosphate-buffered saline (DPBS) on ice. Dissections were completed within 1.5 h.

DPBS was replaced with 250 μL dissociation cocktail containing a mix of 50 μl of dispase (3 mg/ml, Sigma-Aldrich, D4818, 2 mg), 75 μl collagenase I (100 mg/ml, Invitrogen, 17100-017) and 125 μl trypsin-EDTA (0.05%, Invitrogen 25300054). Brains were incubated for 10 min at 25°C with gentle shaking at 75 rpm. Digestion was stopped by adding 500 μL complete Schneider’s medium containing 10% fetal bovine serum. Samples were washed once with complete Schneider’s medium and twice with DPBS. Tissue was then resuspended in 1 mL DPBS containing 1% bovine serum albumin (BSA) and gently dissociated by pipetting with a P1000 until no visible cell clumps remained. The cell suspension was filtered through a cell-strainer cap into a 5 mL BD Falcon FACS tube, and the strainer was washed with an additional 0.5 mL DPBS containing 1% BSA.

Cells were sorted immediately using Melody cell sorter. Sorting was performed using gentle settings with an 85 μm nozzle and 20 psi pressure. GFP-positive cells were isolated from the live singlet population and sorted into 0.5ml DPBS containing 10% BSA.

### Nuclei isolation buffers

1× lysis buffer was prepared with 10 mM Tris-HCl pH 7.4, 10 mM NaCl, 3 mM MgCl₂, 0.1% Tween-20, 0.1% Nonidet P40 Substitute, 0.01% digitonin, and 1% BSA in nuclease-free water. Nonidet P40 Substitute was prepared as a 10% stock when using Sigma 74385. Digitonin was warmed to 65°C as needed to dissolve precipitates before use. For multiome experiment, additional RNase inhibitor was added to a final concentration of 1 U/µl to prevent RNA degradation. Similarly RNase inhibitor was added to the Nuclei Buffer to a final concentration of 1 U/µl as well.

Lysis dilution buffer was prepared with 10 mM Tris-HCl pH 7.4, 10 mM NaCl, 3 mM MgCl₂, and 1% BSA in nuclease-free water. 0.1× lysis buffer was freshly prepared by diluting 1× lysis buffer 1:10 in lysis dilution buffer. For example, 200 μL 1× lysis buffer was mixed with 1.8 mL lysis dilution buffer to prepare 2 mL 0.1× lysis buffer.

### Nuclei preparation, library construction, and sequencing

Single-cell ATAC libraries and the multiome library were prepared and sequenced at the DNA Services laboratory of the Roy J. Carver Biotechnology Center at the University of Illinois at Urbana-Champaign. FACS-sorted cells were immediately concentrated by centrifugation at 500 × g for 10 min at 4°C. The supernatant was carefully removed, and cells were resuspended in chilled 0.1× lysis buffer and incubated on ice for 4 min. Chilled wash buffer was then added to the lysed cells, and nuclei were collected by centrifugation at 500 × g for 10 min at 4°C. The washing step was repeated three times for the multiome experiment. The supernatant was removed without disturbing the nuclei pellet, and nuclei were resuspended in diluted nuclei buffer and delivered to the facility. Nuclei concentration was determined using the Nexcelom K2 brightfield/dual florescence cell counter (Nexcelom Biosciences, Lawrence MA) with AO/PI staining. After nuclei isolation, the viability was checked to be between 1-5% to confirm heathy disruption of the cell membrane. The nuclei were also checked by microscopy to ensure that they maintained a healthy morphology without any blebs or leakages. The nuclei were washed with PBS buffer containing 0.04% BSA, then recounted for library preparation.

For single-nuclei ATAC-seq experiment, the target number of nuclei (10,000) from each population were converted into individually barcoded ATAC DNA libraries with the Chromium Next GEM Single Cell ATAC Reagent Kit V2 from 10X Genomics (Pleasanton, CA) following the manufacturer’s protocols. Briefly, nuclei are incubated in transposition mix 30 min at 37C prior to GEM generation to allow preferential transposition of DNA in open regions of chromatin. After GEM generation, the 10x barcoded DNAs are collected and individually barcoded dual-index libraries compatible with the Illumina chemistry are constructed. The final libraries are quantitated on Qubit (Life Technologies, Grand Island, NY) and the average size determined on the AATI Fragment Analyzer (Agilent Technologies, Santa Clara, CA). Libraries are pooled evenly, and the final pool diluted to 5nM final concentration. The 5nM dilution is further quantitated by qPCR on a BioRad CFX Connect Real-Time System (Bio-Rad Laboratories, Inc. CA).

For single-nucleus multiome experiments, the target number of nuclei (10,000) were processed using the 10x Genomics Chromium Next GEM Single Cell Multiome ATAC + Gene Expression (GEX) workflow according to the manufacturer’s protocol. Gene expression cDNA libraries and ATAC libraries were generated from the same nuclei using Chromium Next GEM Single Cell Multiome ATAC + Gene Expression Reagent Bundle, 4 rxns (Product: 1000285). Similar pre-sequencing preparation was also done for the multiome libraries.

Four single-nucleus ATAC-seq libraries were sequenced on two SP 2×50nt lanes in a NovaSeq 6000 flowcell as paired-reads with 50nt in length. The first and second reads of the ATAC libraries contain the insert gDNA sequence, while the first index read contains the i7 index and the second index reads contains both the i5 index and the 10x barcode information. The ATAC and GEX libraries from the multiome experiment were sequenced after fragment analysis and qPCR quantification, on separate 25B lanes on a NovaSeq X Plus with V1.0 sequencing kits. Read lengths for the GEX llibrary were: Read1 is 28nt in length, Read2 is 150nt in length, I1 and I2 are 10nt in length. Read lengths of the ATAC library were: R1 has length of 50nt, R2 has length of 49nt, UMI barcodes are in I2 reads and have length of 24nt, indices are in I1 reads and have length of 8nt. The read lengths were set as recomended by 10xgenomics.

### Building single-cell assay datasets

1.6 Billion and 1.9 Billion read pairs were obtained for ATAC and GEX libraries respectively in the multiome experiment. cellranger-arc-2.0.2 from 10x Genomics was used with *Drosophila melanogaster* BDGP6.32.109 reference genome and filtered reference *Drosophila melanogaster* BDGP6.32.109.gtf file for alignment and downstream dataset building. cellranger-arc count was used to call cells and build the single-cell multiome dataset for further downstream analysis. From the 4 sc-ATAC libraries processed, a total of 1.8 Billion read pairs were obtained. The sc-ATAC library was similarly built following the cellranger-atac/2.0.0 pipeline. There were four libraries parallelly produced and each were aligned to the BDGP6.41 *Drosophila* reference genome. Basecalling and demultiplexing of raw data were done with the mkfastq command of the software Cell Ranger-ATAC (10x Genomics). The 4 ATAC libraries aggregated to a total of 44,463 cell barcodes and the multiome experiment gave 7,358 cell barcodes for downstream analysis.

### Single-cell data analysis

All downstream analyses were performed in R version 4.5.2. Quality control and single-cell analyses were performed using Seurat version 5.3.1 and Signac version 1.16.0. scATAC-seq and multiome datasets were analyzed following the Signac workflow with default parameters unless otherwise specified.

For multiome data, cells were retained if they met the following quality-control criteria: ATAC counts between 100 and 50,000 per cell, RNA counts between 100 and 15,000 UMIs per cell, nucleosome signal <10, blacklist ratio <0.05, and mitochondrial RNA percentage <15%. Dimensional reduction, clustering, integration, and visualization were performed using standard Seurat and Signac functions. Integration of scATAC-seq and multiome datasets was performed according to the Signac integration workflow.

Clusters were annotated based on published medulla single-cell datasets and known marker expression. Feature plots, coverage plots, and peak-gene linkage analyses were generated using Seurat and Signac. LinkPeaks function of Signac was used to link accessible regions with gene expression of corresponding genes. Candidate regulatory elements were selected based on chromatin accessibility, peak-gene linkage, nearby nTF expression, and correspondence with cell-type- or Notch-status-specific expression patterns. RunChromVAR function of Signac was used to find deviations between chromatin accessibility and motif enrichment at peak regions for different transcription factors. Motifs were obtained from JASPAR2024 database. Format adjusted BSgenome.Dmelanogaster.UCSC.dm6 reference assembly was used for these analyses.

### Reporter construct generation

Candidate enhancer fragments were PCR-amplified from yw genomic DNA using region-specific primers. Mutated enhancer sequences were synthesized as gBlocks by Integrated DNA Technologies. Enhancer fragments were cloned into the pJR12-GFP::PEST reporter vector upstream of the hsp70 minimal promoter using NotI and AscI restriction sites.

The enhancer regions cloned in this study were:

bsh#4: 2L:19,772,130–19,772,979
bsh#5: 2L:19,773,292–19,774,105
TfAP-2#3: 3L:21,587,587–21,588,527
tup#14: 2L:18,884,361–18,885,322
tup#20: 2L:18,892,536–18,893,464
Lim3#13: 2L:19,097,835–19,099,212

### Generation of transgenic reporter flies

Reporter transgenes were inserted into the VK00027 landing site on the third chromosome (BDSC 35569) by φC31 integrase-mediated transgenesis. Using the same genomic landing site minimized position effects among reporter constructs. Red-eyed male transformants were recovered and crossed individually to establish stable transgenic fly lines. Reporter expression was examined in third-instar larval brains or adult brains by immunostaining with anti-GFP and appropriate cell-type marker antibodies.

### Enhancer deletion

Endogenous enhancer deletions were generated using a CRISPR/Cas9-mediated precise genome editing strategy (Ref). Guide RNAs flanking the target enhancer interval were designed and cloned into gRNA expression vectors, and donor constructs were generated for homology-directed repair. Transgenic flies carrying candidate deletion alleles were screened based on red fluorescence in the eye. The reporter cassette was subsequently removed by crossing to αTub-piggyBacK10 (BDSC 32070). Deletions were validated by sequencing.

For deletion of the bsh#4/#5 region(2L:19,772,106–19,774,194), the following gRNAs were used: gRNA1, AACTGGGTAATCTTCAATAG; gRNA2, ATTTGATTGCGAGTTACGTG; gRNA3, TCGACTATAGCAACCCATCT; and gRNA4, GCGCTTCACTATTTGTGTAA.

### Immunostaining

Third-instar larval brains or adult brains were dissected in 1× PBS and fixed in 4% formaldehyde on ice for 30 min. Samples were washed in PBST and incubated with primary antibodies using standard immunostaining procedures.

Primary antibodies generously provided by Dr. Claude Desplan included anti-Bsh (rabbit, 1:500), anti-TfAP-2 (guinea pig, 1:100), anti-Erm (rat, 1:200), anti-Run (guinea pig, 1:500), anti-Lim3 (guinea pig, 1:500), anti-Ap (rat, 1:100), anti-Vvl (guinea pig, 1:250), anti-Scro (guinea pig, 1:100), anti-Toy (rabbit, 1:500), anti-Sox102F (rabbit, 1:200), anti-Hbn (rabbit, 1:500), and anti-Oc (guinea pig, 1:500). Anti-Hey antibody was provided by Dr. Christos Delidakis (guinea pig, 1:1000), and anti-Sim antibody was provided by Dr. Brian Gebelein (guinea pig, 1:500).

Commercial antibodies included anti-Tup (DSHB, mouse 40.3A4, 1:50), anti-Dac (DSHB, mouse mAbdac2-3, 1:20), anti-GFP (AbD Serotec, 4745-1051, sheep, 1:500), and anti-RFP (Abcam, ab62341, rabbit, 1:1000). Secondary antibodies were obtained from Jackson ImmunoResearch or Invitrogen.

### HCR RNA-FISH

HCR RNA-FISH probes against Lim3 and tap were obtained from Molecular Instruments. A published protocol^47^ was followed. Third-instar larval brains were dissected in cold 1× PBS and fixed in 4% formaldehyde on ice for 30 min. Samples were rinsed three times with PBST containing 1% Triton X-100 and washed four times for 5 min each at room temperature.

Samples were pre-hybridized in 200 μL pre-warmed probe hybridization buffer for 30 min at 37°C. Probe solution was prepared by adding 2 μL of each HCR HiFi probe to 100 μL HCR probe hybridization buffer. Samples were incubated in probe solution overnight for 12–16 h at 37°C. The next day, probe solution was removed, and samples were washed four times for 15 min each at 37°C with pre-warmed probe wash buffer. Samples were then washed three times for 5 min each at room temperature in 5× SSCT.

Hairpin amplification was performed by replacing the pre-amplification solution with hairpin solution and incubating samples overnight for 12–16 h at room temperature. On the third day, hairpin solution was removed, and samples were washed in 5× SSCT twice for 5 min, twice for 30 min, and once for 5 min at room temperature. Immunostaining was performed after HCR RNA-FISH when required.

### Imaging and image processing

Confocal images were acquired using Zeiss LSM 700 or Zeiss LSM 900 confocal microscopes. Images were processed using Adobe Photoshop and assembled using Adobe Illustrator. Equivalent imaging and processing conditions were used for comparisons within each experiment.

### Statistics

Ap intensity was quantified in Fiji. Ap intensity in RNAi clones was normalized to adjacent wild-type tissue from the same brain. Each point represents one brain, with at least two clones measured per brain; five brains were analyzed in total. The dashed line indicates WT-normalized intensity = 1. Mean ± SEM is shown. Significance was assessed by one-sample t test against 1.

## Data availability

Raw and processed sequencing data will be deposited in the Gene Expression Omnibus and will be available upon publication.

## Code availability

Code used for data processing and analysis will be made available upon publication or upon reasonable request.

## Acknowledgments

We thank the Flow Cytometry Facility, the DNA Services Laboratory, and the High-Performance Computing in Biology Group at the Roy J. Carver Biotechnology Center, University of Illinois Urbana-Champaign, for assistance with FACS sorting, 10x single-cell library construction, sequencing, and computational support. We thank the fly community, especially Dr. Claude Desplan, Dr. Christos Delidakis, and Dr. Brian Gebelein, for generously providing antibodies and fly stocks. We also thank the Bloomington Drosophila Stock Center, FlyORF, the Vienna Drosophila RNAi Center, the Developmental Studies Hybridoma Bank, and the TRiP at Harvard Medical School, for fly stocks and reagents. We thank members of the Li lab for helpful discussions. This work was supported by the National Eye Institute grant R01 EY026965-01A1 and 2 R01 EY026965-06A0 to X.L.

## Author contributions

Conceptualization: X.L. and Y.Z.; Methodology: Y.Z., T.S. H.Z. and X.L.; Software: Y.Z., and T.S.; Validation: Y.Z. and T.S.; Formal Analysis: Y.Z. and T. S.; Investigation: Y.Z., T.S. H.Z, R.W.J, and M.S.; Data Curation: Y.Z. and T.S., Writing - Original Draft: Y.Z., T.S. and X.L, Writing - Review & Editing: all authors, Visualization: Y.Z., Supervision: X.L. and Y.Z., Project Administration: X.L., Funding Acquisition: X.L..

---

**Figure S1.**
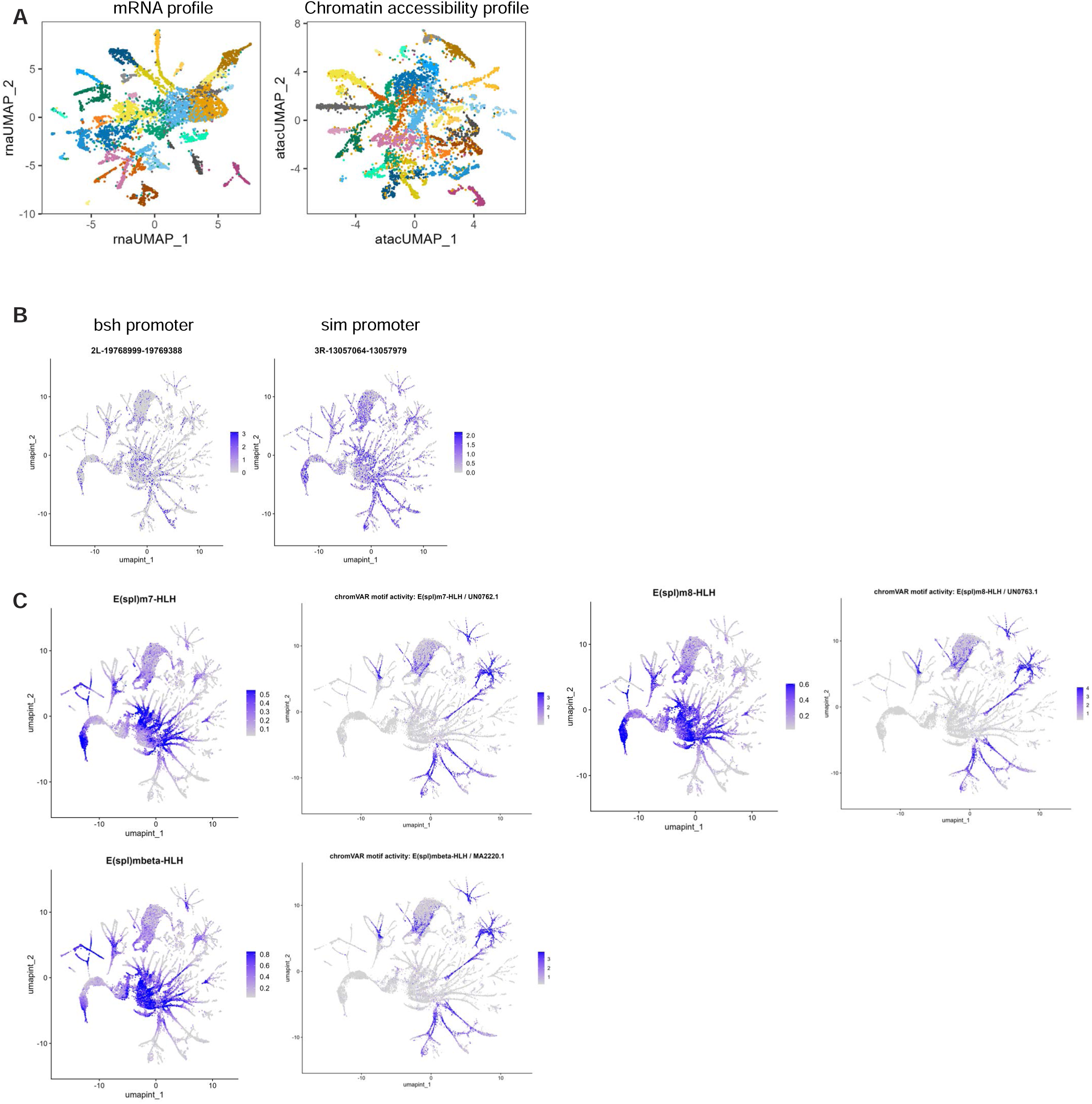
Multiome embeddings, promoter accessibility, and bHLH motif activity. (A) UMAP visualization of the single-cell multiome dataset generated using either the RNA modality or the ATAC modality, showing the corresponding transcriptomic and chromatin accessibility profiles. (B) Feature plots showing accessibility at representative nTF promoters, including the bsh and sim promoters. Promoter accessibility is broadly distributed across cell states and shows limited cell-type or Notch-status specificity. (C) Feature plots showing the expression of representative E(spl)/Hey-family bHLH factors and chromVAR deviation scores for their corresponding motifs.

**Figure S2.**
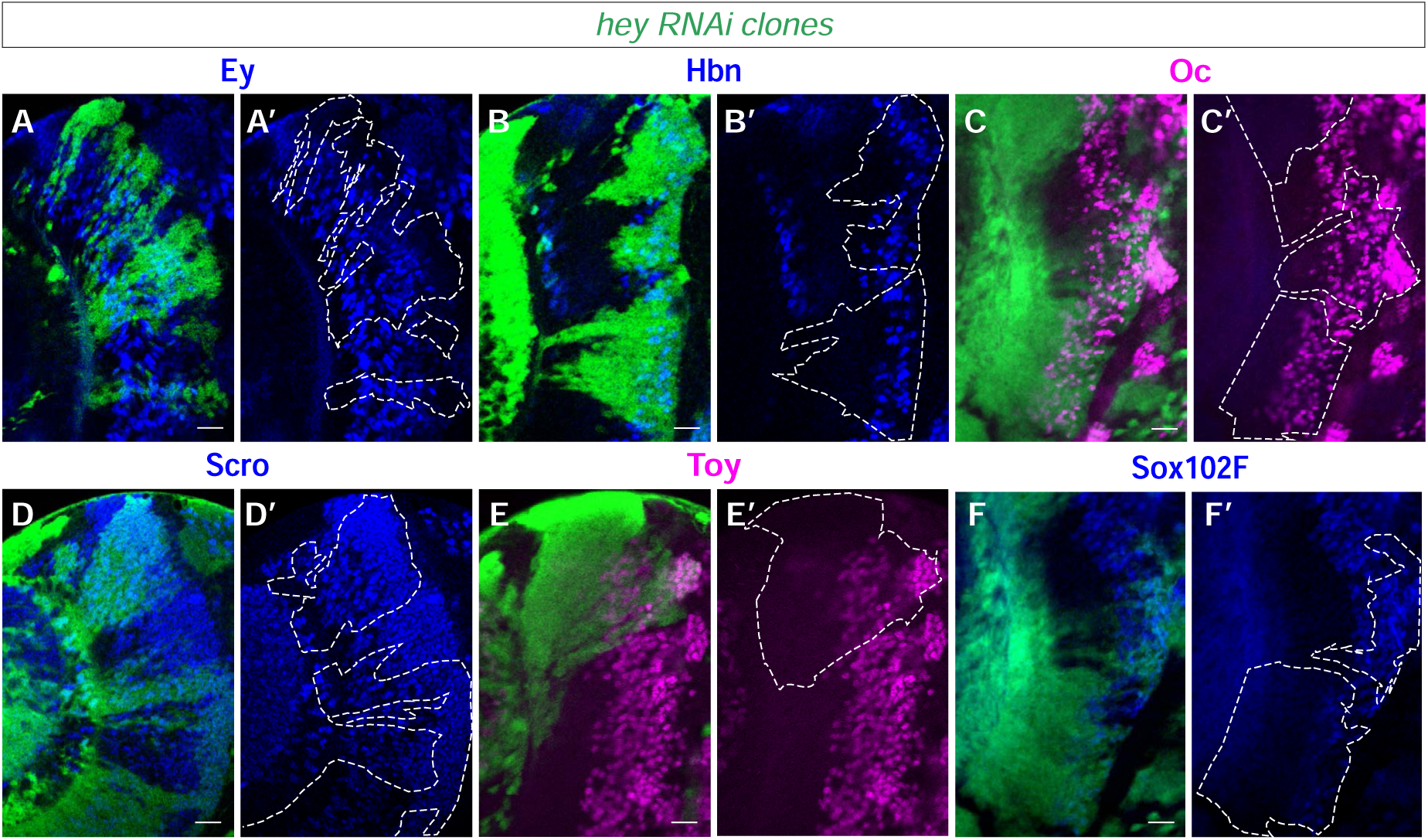
Additional Notch-off and Notch-on nTFs are not ectopically expanded after *Hey* depletion. (A–C) Notch-off nTF expression is not ectopically expanded in *Hey* RNAi clones. Ey (A–A′), Hbn (B–B′), and Oc (C–C′) remain largely restricted outside clone regions. (D–F) Additional Notch-on nTFs are largely preserved in *Hey* RNAi clones. Scro (D–D′), Toy (E–E′), and Sox102F (F–F′) expression remain detectable in clone regions. White dashed outlines mark *Hey* RNAi clone regions. Scale bars, 20 μm.

**Figure S3.**
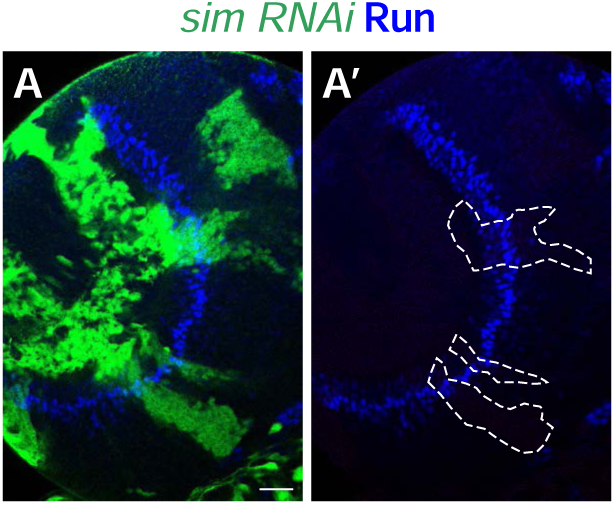
*sim* RNAi does not alter Run expression in Mi4 neurons. (A and A′) Run expression is preserved in *sim* RNAi clones. Clone regions are marked by GFP and outlined with white dashed lines in the single-channel view (A′), indicating that loss of sim does not transform Notch-on TfAP-2+ cells into Run+ Notch-off sibling identity. White dashed outlines mark *sim* RNAi clone regions. Scale bar, 20 μm.

**Figure S4.**
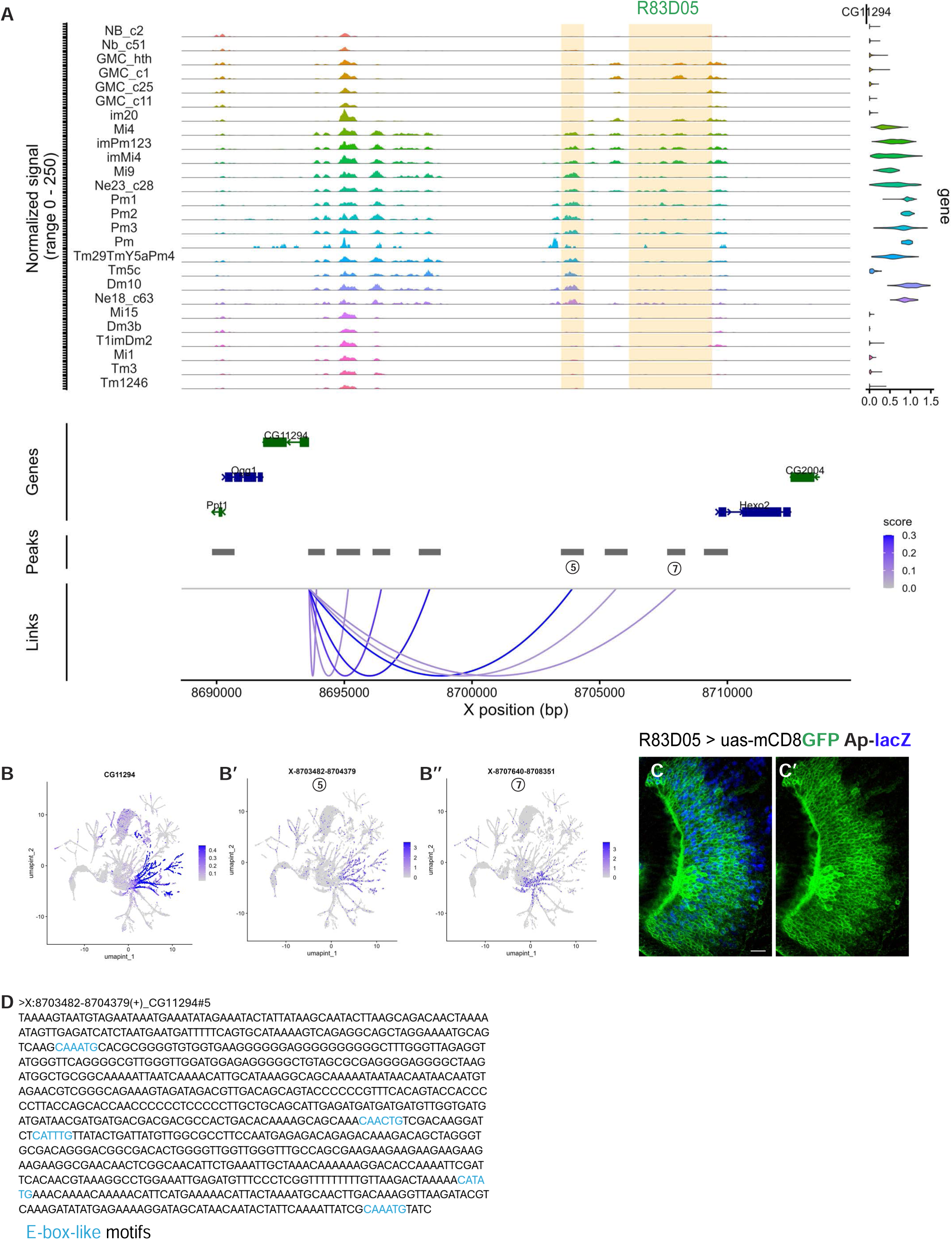
Candidate CG11294 regulatory elements suggest distributed temporal and Notch-status-associated enhancer logic. (A) Genome browser coverage plot showing chromatin accessibility and peak-gene links at the CG11294 locus across developing medulla cell clusters. Candidate linked elements CG11294#5 and CG11294#7 are indicated. The Janelia GAL4 fragment R83D05, which spans the CG11294#7 region, is highlighted. (B–B″) Feature plots showing CG11294 expression (B) and accessibility of candidate linked elements CG11294#5 (B′) and CG11294#7 (B″). CG11294#7 shows a temporal accessibility pattern, whereas CG11294#5 represents a candidate Notch-off-associated element. (C and C′) R83D05 enhancer activity in larval optic lobes. R83D05 drives reporter expression within the CG11294-associated temporal domain and overlaps with ap-lacZ+ Notch-on cells, suggesting that this regulatory region encodes temporal information but does not fully restrict expression to the endogenous Notch-off CG11294 pattern. (D) Sequence of the CG11294#5 element. Predicted E-box-like motifs are highlighted. Scale bars, 20 μm.

## Description of Supplementary Tables

**Supplementary Table 1. Candidate cis-regulatory elements linked to gene expression programs.**

Summary of candidate regulatory elements identified by single-cell multiome peak-gene linkage analysis. For each gene, the table lists the linked accessible peak, genomic coordinates, linkage score, z-score, and p value.

**Supplementary Table 2. Motifs enriched in chromatin regions preferentially accessible in Notch-on neurons.**

Summary of motif enrichment analysis comparing differentially accessible chromatin regions in Notch-on neurons with those in Notch-off neurons. For each motif, the table lists the motif ID, number and percentage of Notch-on-enriched regions containing the motif, background frequency, fold enrichment, p value, adjusted p value, and motif name.

